# Microvascular network organization and hemodynamic perfusion protect the brain against hypoxia

**DOI:** 10.64898/2026.02.03.703617

**Authors:** Thomas Ventimiglia, Frédéric Lesage, Andreas A. Linninger

**Affiliations:** Department of Biomedical Engineering, University of Illinois, Chicago, USA; Department of Electrical Engineering, Polytechnique Montreal, Canada; Department of Neurosurgery, University of Illinois, Chicago, USA

**Keywords:** hemodynamics, neurovascular unit, capillaries, imaging, neuropathology

## Abstract

Mechanistic simulations of blood flow and oxygen exchange showed regions of cortical tissue tolerating substantial increase in local oxygen consumption (CMRO2) before reaching hypoxia (pO2<10 mmHg). The observed robustness in O_2_ supply was attributed to overcapacity in convective oxygen transport in the pial arterial network combined with a surplus in the number of capillary flow paths. Microcirculatory flux analysis suggests that *network* induced hemodynamic flow patterns impart *intrinsic reserve* to protect the brain against perfusion variances or metabolic demand surges during activation. Furthermore, oxygen transport in cortical tissue is characterized by two regimes: in the *transport zone —* centered on *penetrating arteriole trees* composed of a single penetrating vessel connected to the post-arteriole capillary transition zone — strong diffusion supports high oxygen tension with only modest contribution from capillaries. This regime transitions into the *terminal/reactive zone* where oxygenation is sensitive to capillary density and perfusion. Quasi-dynamic simulations also enabled reconstruction of the BOLD signal underlying functional imaging. Simulations at single micron resolution further show that age-related reductions in arterial saturation and systemic hematocrit were sufficient to induce hypoxic tissue pockets in the terminal zone at nominal perfusion (CBF) and metabolic activity (CMRO_2_), and neutrophil adhesion induced capillary flow stalling further exacerbates hypoxia.

## Introduction

Microcirculatory events^1,2^ or hypoperfusion in ischemic stroke^3,4^ have been implicated in certain neurovascular pathologies. A study in rodent AD model showed that vascular inflammation led to the adhesion of neutrophils in capillaries, effectively reducing perfusion by blocking red blood cell passage.^5^ In a separate study, aged mice populations had a decrease in tissue oxygen accompanied by reduced hematocrit and the rare emergence of hypoxic micro-pockets acquired by two photon phosphorescence O_2_ microimaging.^2^ While both observations could be related by the fact that RBCs carry oxygen, specific mechanisms of age or hypoperfusion related causes and theoretical verification of microscale disruptions to neurovascular coupling are missing. The classical Krogh’s cylinder^6^ or the Crone-Renkin model^7,8^ illuminate solute exchange around a single capillary; but their assumptions are often invalid in pathologies nor do they account for microvascular network effects.

Mechanistic modeling of blood flow in anatomically realistic network geometries aim to predict hemodynamics and solute exchange without the need for restrictive averaging assumptions. Image-based reconstructions of vascular graphs^9–15^ faithfully replicate key network properties such as connectivity, local capillary length density, and bifurcation-to-bifurcation and tissue-to-vessel distances.^16,17^ Moreover, detailed biphasic blood flow models sensitive to critical perfusion parameters such as CBF, CBV, hematocrit, as well as shear and hematocrit dependent blood rheology^10^ are able to quantify complex network effects including plasma skimming^10,18^ and graded oxygen extraction along penetrating arteries and veins.^19^

A critical modeling element concerns blood-tissue solute exchange^9–11,20,21^ to account for O_2_-bound convection in blood, dissociation from hemoglobin, solute transfer across the vascular wall, diffusion to neurons and glia and finally oxygen consumption by cell metabolism (CMRO_2_). Because the characteristic length of micro-transport phenomena is in the order of capillary diameters (and individual cells, L∼2-5 µm), space discretization needs a very fine-grained resolution of 1 µm or less for reliable predictions.

### Dual-domain technique

Simulating solute exchange at the micron scale is intractable for standard numerical techniques. We specifically designed a numerical approach entitled *dual-domain technique*^9,20^ for predicting O_2_ exchange in capillary beds down to single micron resolution.^21,22^ The dual-domain technique has sufficient resolution to quantify transport between cell clusters and individual capillaries. The use of transport theory is innovative, because current metabolic imaging does not yet have sufficiently high spatial resolution nor can raw image data explain the underlying mechanisms of solute exchange in the brain. The current study uses ultra-high resolution simulations to expound physicochemical principles that underpin the effects of local disruptions to O_2_ exchange caused by age-related leucocyte stalling or micro-strokes.

### Outline

We performed mechanistic simulations of O_2_ convection in blood, extraction, and metabolic turnover in the murine cortex. Micron-scale oxygen flux analysis was used to elucidate mechanisms preventing tissue from under-perfusion. Conditions provoking hypoxia in cortical tissue were explored by systematically modulating local CBF and CMRO_2_. Finally, we compared simulated hypoxic regions to in-vivo experiments.^23^ We predicted and validated frequency, size and location of hypoxic micro-pockets^2^ under simulated disturbances modelled on age-related changes.

## Methods

### Representation of neurovascular anatomy

The vascular anatomical network (VAN) is represented as a graph of nodes (vertices) and faces (edges). Cortical micro-domains consisting of tissue and blood vessels are registered to a 3D-array of cubic voxels in natural (Cartesian) coordinates as shown in Fig. 1. Each vascular segment is digitally represented as a cylindrical tube of given length and diameter (Fig. 1, panels A-D). Registration of the vascular graph to the 3D voxel array is achieved by constructing a cylindrical surface with spherical endcaps around each vessel: each voxel of the 3D array is then labelled as either belonging to the interior lumen of the blood vessel (red), an interface with solute exchange between blood and tissue (endothelial wall, gray), or extravascular tissue (blue in Fig. 1, panels E-I). Implementation details on the registration of the vascular graph to the 3D voxel array are discussed in supplement section 1.

**Figure 1.**
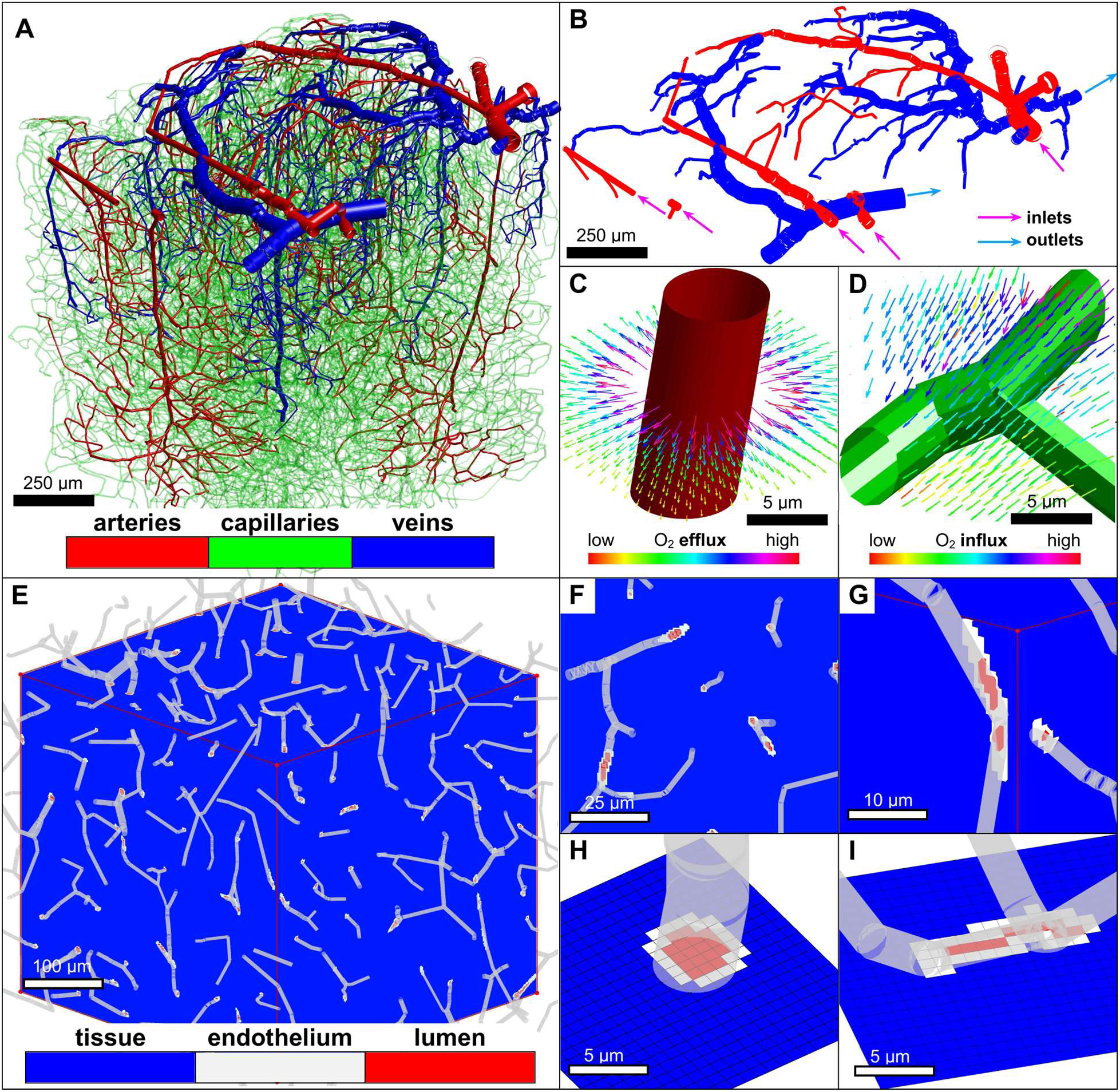
Overview of discretized computational domains derived from VAN. A) In-vivo mouse cortical network derived from two-photon imaging^16^. B) Pial surface vessels with inflow and outflow boundaries indicated. C,D) Thin penetrating arteriole and capillary bifurcation with surrounding oxygen flux illustrated by arrows. The shown capillary junction is an example of oxygen flowing from tissue back into the vascular compartment. E) Cartesian voxel array representation of coupled network and tissue. F,G) Detail of Cartesian voxel array with labelled voxels for extravascular tissue, endothelial tissue, and interior vessel lumen. H,I) Detail of Cartesian voxel registration of a penetrating arteriole and capillaries respectively.

### Hemodynamics

Blood flow is modeled as a biphasic suspension, as described in prior works.^10,18,19^ Mathematical details are found in supplement sections 2 and 3. We performed simulations in a 3×3×1 mm^3^ cut-out curated from a micro-CT study (photomicrographs)^17^ of a whole mouse brain connectome and a smaller 1×1×1 mm^3^ in-vivo network obtained by two-photon imaging.^11,16^ The whole brain cutout is shown in context along with simulated vascular pO_2_, hematocrit, and flow fields in Fig. 3 panels A-D respectively. We also repeated simulations on a 3×3×1.2 mm^3^ synthetic blood flow network whose vascular topology matches in-vivo mouse data (‘digital twins’).^20,24^ We chose extended domain dimensions (3 mm range), so that the core of the predicted O_2_ field (confined to 1×1×0.8 mm^3^ central region) was undisturbed by boundary effects which arise due to severed penetrating arterial branches and capillary segments, causing unnaturally low O_2_ tension and perfusion in border zones. Analysis of oxygen fields for the simulated networks were performed in this 1 mm side-length core. All trends were observed and reproduced in both the anatomical and the synthetic networks.

### Mouse data

Comparison with in-vivo data on hypoxia were based on experiments in young and aged mouse population,^2^ with more details in section 5 of the supplemental material.

### Solute transport and exchange

A central piece of the mathematical model for metabolic neurovascular coupling concerns solute exchange, which entails multiple transport and chemical reaction processes in both blood and tissue coupled by mass transfer between the two domains. In the vascular compartment, oxygen enters with the blood stream with convective blood flow fluxes computed as in supplement section 2. Hemoglobin-bound oxygen enters the microcirculation at arterial inlets (Fig. 1, panel B) with a prescribed saturation. Bound oxygen concentration **c***_v_* is able to dissociate from RBCs using the well-known Hill equilibrium (supplement section 2) becoming free O_2_: *K*(**c***_v_*). Mass transfer into the tissue is modeled as a passive process driven by the concentration difference between O_2_ in blood, *K*(**c***_v_*) and tissue, **c***_t_*. In tissue, extravasated oxygen travels to cells by diffusion, but is also consumed in metabolic reactions to supply neural and glial cell populations. No-flux boundary conditions are enforced at the limits of the rectangular domain. To eliminate boundary effects on predicted oxygen trends, we solve over very large tissue domains (3×3×1.2 mm^3^), but limit our analysis to a central core ROI (1×1×0.8 mm^3^)^20^ undisturbed by far-away domain borders (at least 1 mm distance to border).

### Mass transfer coupling

By registering the vascular graph to the tissue voxel array (supplement section 1), we construct a bipartite graph connecting the nodes of the vascular domain (a directed acyclic graph) to voxels of the tissue domain (graph of the 3D Cartesian tissue domain). The vascular mesh (network graph) is anchored to tissue cells (Cartesian voxel array) at voxels corresponding to the endothelial wall: vascular oxygen is transferred from a given graph node to the endothelial voxels closest to it. The graph theoretical aspects of this problem formulation are explained in more detail elsewhere.^21,22^

### Equation for multi-domain solute exchange

The coupled vascular-tissue O_2_ concentration field at steady-state is expressed in matrix form by equation (1) with more details in supplement section 2. Here, **M**(**f**) is the convection matrix associated with the computed flow field, **V***_t_* is the diagonal matrix of tissue voxel volumes, *K* is the nonlinear Hill operator, and *R* is the Michaelis-Menten reaction rate defined in equation (2). The matrix **C Γ C**^T^ is the *weighted Laplacian matrix* for the coupled vascular and tissue graph; **C** is the incidence matrix and **Γ** is the diagonal matrix of oxygen transport conductances across an edge. The right-hand side is a source term for steady inflow of arterial oxygen at the prescribed concentration ***c̅***_v_. The term **Q**_in_(**f**) accounts for oxygen inflow boundary conditions.

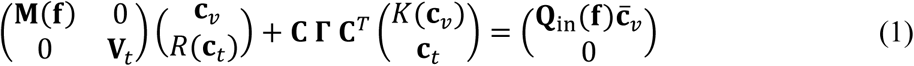

Edge conductances are set as follows: (i) Conductance between tissue nodes is *h*Γ*_t_*, for tissue diffusivity Γ*_t_*, and voxel side length *h*. (ii) Between vascular nodes conductance is zero, following the observation that diffusion of intravascular oxygen in the blood stream is negligible compared to convection.^25^ (iii) For mass transfer between vascular and tissue domains, the conductance is given by *UA*/*w*, where *U* is the permeability of the vessel wall, *A* is the surface area of the vessel wall through which mass transfer occurs, and *w* is the thickness of the endothelial wall.

### Kinetics

O_2_-dissociation from HbO_2_ bound to free (unbound) O_2_ in plasma is modeled via the non-linear Hill equation^26^ shown in supplement section 2. The metabolic reaction rate of oxygen in tissues is modeled via the Michaelis-Menten type rate equation (3):

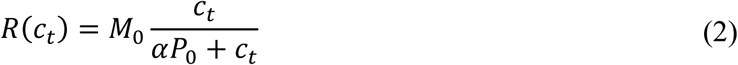

### Newton iteration

Equation (1) is nonlinear due to Hill dissociation *K*(**c***_v_*) and Michaelis-Menten reaction rates *R*(**c***_t_*); hence it is solved iteratively by Newton’s method with details given in supplement section 3. Parameter values for the simulations are given in supplementary Table S1. Choices of nominal conditions are listed in supplement section 4.

## Results

### Uniform extravasation and reabsorption across vessel types

The results of Fig. 1. were generated for an in-vivo 1×1×1 mm^3^ vascular network extracted from the somatosensory cortex and imaged with two-photon microscopy.^16^ Oxygen fluxes in the tissue domain are visualized as arrows, showing that directionality of O_2_ flows is mostly extravasating. In many instances, oxygen may transfer back from the tissue space into blood in some segments of the network; Fig. 1 panel D, shows such an example in a capillary deep in the cortex and far downstream of the nearest pial inlet.

Simulations for nominal conditions were also carried out on the 3×3×1.2 mm^3^ synthetic network^20,24^ with analysis limited to the 1×1×0.8 mm^3^ core region. The pdfs of Fig. 2, panels A-C show that capillaries, arterioles, and venules all exhibit extravasation, but there are also segments with some reabsorption. The net O_2_ transfer over larger control volumes (e.g. side length 250 µm) is always positive (extravasating) and nearly constant (Fig. 2, panel D). Net extravasation per unit surface area is similar in capillaries and arterioles, (on the order of 0.02 µmol/cm^2^/min); but somewhat lower in venules (mean value of 0.01 µmol/cm^2^/min) showing that all vessel types contribute significantly to tissue oxygenation (Fig. 2, panels A-C, and E).

**Figure 2.**
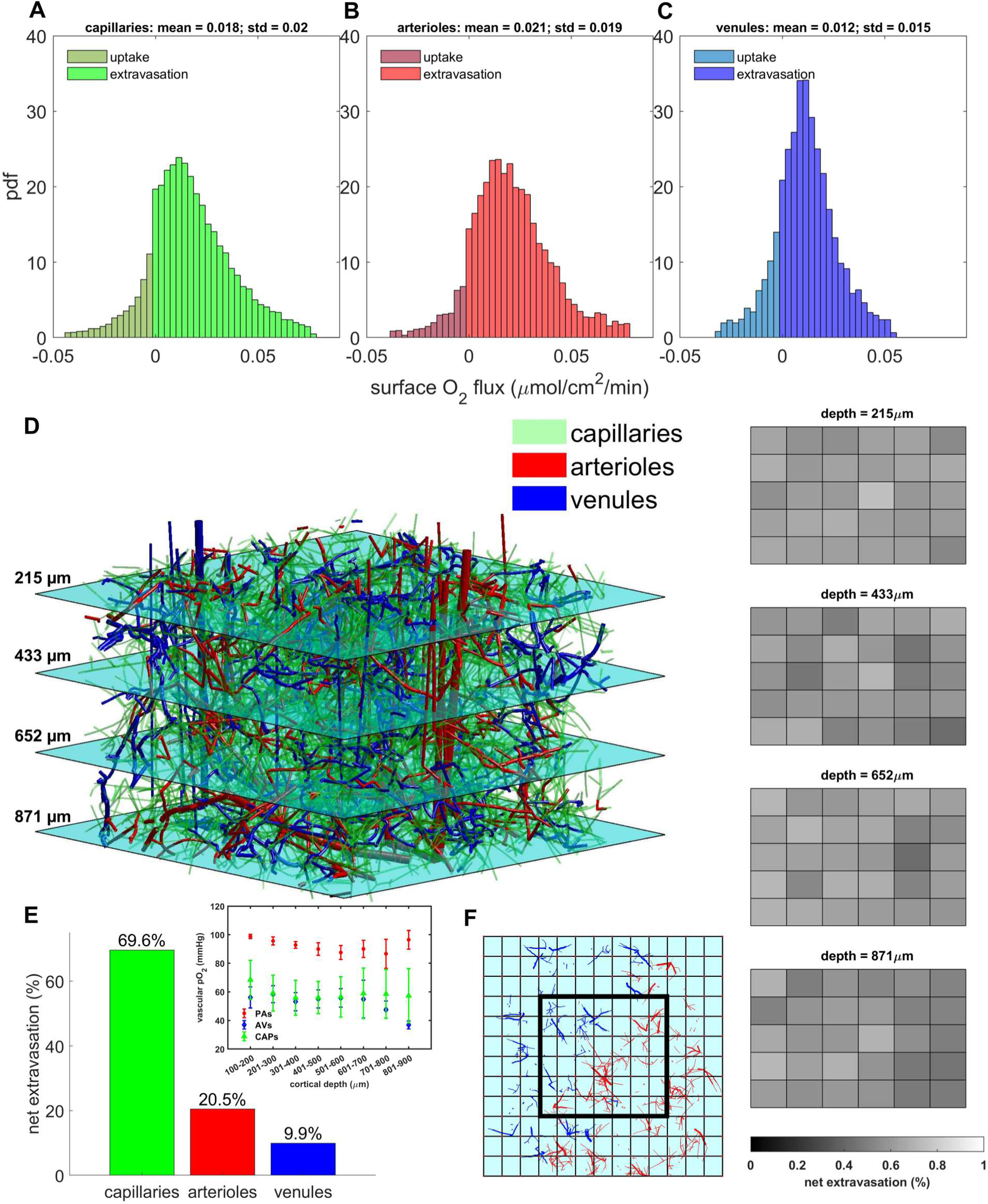
Extravasation flux throughout the 1×1×0.8 mm^3^ core of the 3×3×1 mm^3^ synthetic network in the nominal case. A-C) Extravasation flux over vessel surface areas is approximately equal on average between the capillaries and arterioles. Mean flux from venules is slightly lower owing to their lower intravascular oxygen tension. Negative values indicate uptake of oxygen from tissue back into the vascular compartment. D) Net extravasation in the core zone is divided nearly evenly over coarse cubic voxels of side length 250µm. Slices are shown at different depth levels throughout the core. E) Contribution of capillaries, arterioles, and venules to extravasation show that all three vessel types contribute significantly to oxygen extraction. Inset shows pO_2_ levels with respect to cortical depth in penetrating arterioles (PAs), ascending venules (AVs), and capillaries (CAPs). Bars are mean plus/minus standard deviation of vascular pO_2_ in each 100 µm thick layer. F) Overhead view of core region of interest (thick black outline) over coarse voxels. Red and blue vessels are arterioles and venules respectively.

### Flow stalling in capillaries causes redirection of flow far from the occlusion site

Cruz Hernandez et. al.^5^ reported that neutrophil adhesion can cause stagnation or complete cessation of flow in affected capillary segments. We tested the hypothesis that neutrophil adhesion^5^ suppresses flow in affected capillary segments and that local stalls can induce hypoxic pockets in cortical tissue as observed in Moeini et. al.^2^ Irrespective of the multiple underlying biological aspects of neutrophil adhesion such as neuro-inflammation, we chose to quantify the effect of stalls on micro-perfusion by occluding capillary blood flow in a 3×3×1 mm^3^ in-vivo network. To mimic the hemodynamic effect of stalls, we increased the resistance of all capillary segments (faces) within a spherical region (increasing resistance by a factor of 10^4^ reduces flow in these segments to less than 1% of their nominal value).

Complete capillary stalling within a volume (affecting up to N=2519 capillary segments simultaneously whose center points are within sphere of radius R) reduces overall bulk flow as expected, but also reversed blood flow directions or even increased it elsewhere in the network (Fig. 3, panels R and S). Although blood supply is almost completely stopped in the occluded volume, (reported as a negative flow change in Fig. 3 R, S), local stalling rearranges the entire capillary flow field affecting flow magnitude and direction up to 500 µm from the obstruction site; some flows were more than doubled compared to pre-stalling conditions. These observations are consistent with results reported by Jamshidi et. al.^18^, which had previously showed that capillary stalls reversed flows in significant sections of the capillary bed.

**Figure 3.**
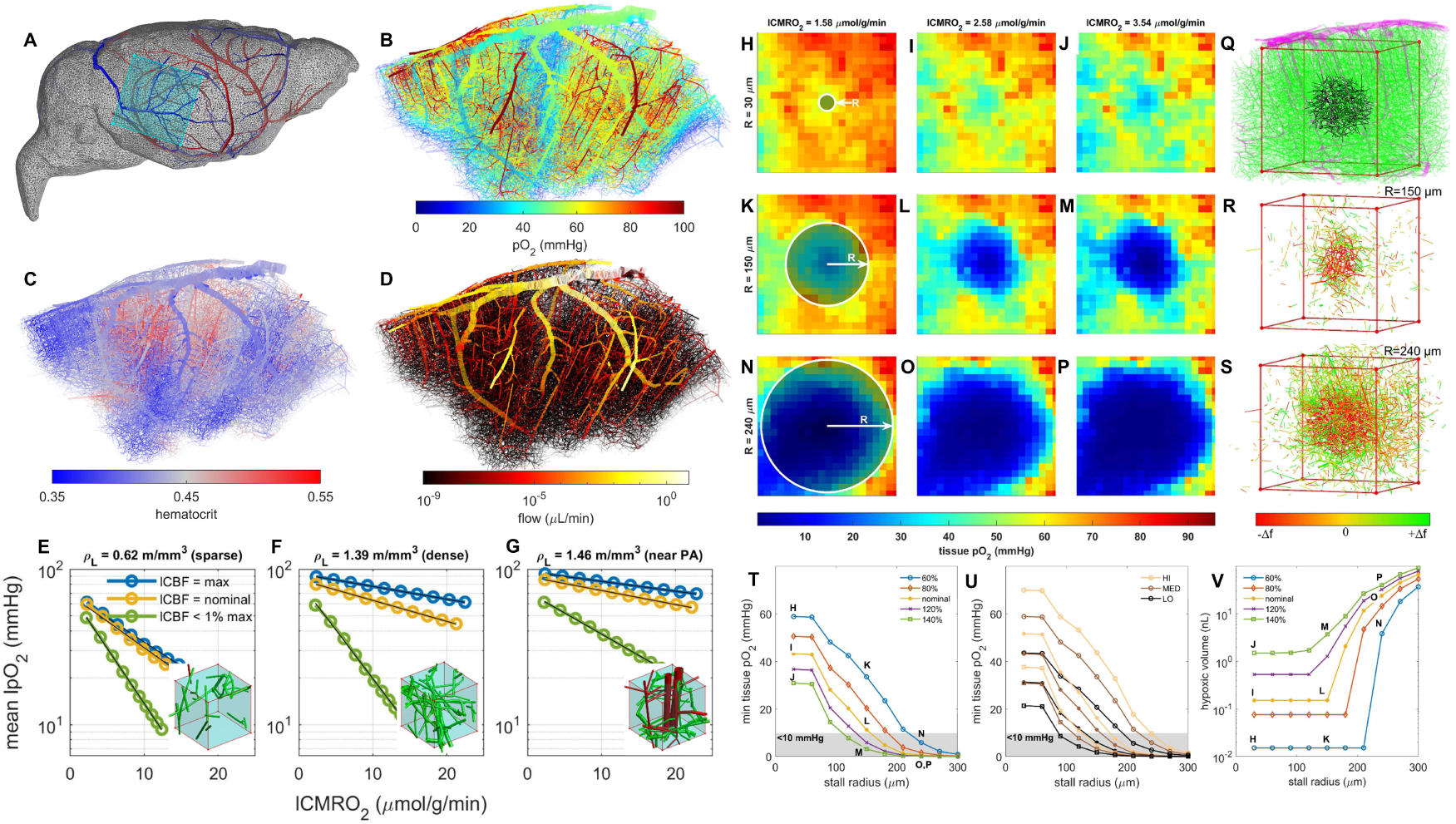
The relation of capillary stalling and density on hypoxia. A) Context view of 3×3×1 mm^3^ cutout from the whole mouse brain data set.^17^ Major veins (blue) and arteries (red) are shown. B-D) Simulations of vascular pO_2_, hematocrit, and blood flow respectively in 3×3×1 mm^3^ cutout. E-G) Mean local tissue pO_2_ with respect to increasing local CMRO_2_ in three ROIs of increasing length density *ρ*_L_. Each trend-line represents distinct CBF levels: the maximum CBF attainable by reducing flow resistance locally, the nominal case, and <1% of the maximum CBF. Insets show microvasculature in the regions corresponding to the simulations in panels E-G respectively: green = capillaries, red = penetrating arteriole. H-P) Cross sections of tissue pO_2_ under different stalling and demand scenarios: these simulations were carried out on a 3×3×1 mm^3^ in-vivo VAN curated by the Kleinfeld lab^17^ and shown in panels A-D. Capillary stalls are confined to spherical regions of increasing radius R delineated by the white outline (displays along rows). We also varied CMRO_2_ over a central cubic ROI with dimensions 1 x 1 x 0.8 mm^3^ (varied along columns). Hypoxia forms at the center of the stalling zone for large stall R and elevated CMRO_2_ indicative of network resilience of the O_2_ supply to stalling events. Q) ROI and spherical stalling zone in context with surrounding network. Magenta vessels are large diameter pial and penetrating vessels. Green vessels are capillaries, which are defined here as having a diameter < 7 µm (Ji. et. al.^17^). Black vessels indicate stalled capillaries within sphere of radius 240 µm. R-S) Vessels with redirected flow around the central stalling zone of radius R for panels K-M and N-P respectively. Redirected flow is extensive throughout the network, far from the central focus of the stalls. T-U) Minimum tissue pO_2_ at the center of the stalling zone and V) volume of hypoxic tissue with respect to stalling extent and reaction rate. The corresponding scenarios in panels H-P are indicated next to the appropriate data point in panels T and V.

### Size of stalling volume required to induce hypoxia depends on lCMRO_2_, hematocrit, and arterial oxygen saturation

To determine the spatial extent necessary to induce hypoxia, capillary stalls were induced in concentric spherical regions with step-wise increment in radius, R: see Fig. 3. We see that the critical radius for which hypoxia forms in the center of the stalling region depends on the local rate of oxygen consumption, as well as on the systemic hematocrit (Fig. 3, panels T, U). For very low demand in the core region, (local CMRO_2_ = 1.58 µmol/g/min and normal CBF = 0.79 mL/g/min, Fig. 3, panels H, K, and N) even complete capillary stalling in spheres with R ≤ 200 µm did not induce hypoxia (Fig. 3, panel T). For nominal reaction rates, (local CMRO_2_ = 2.58 µmol/g/min, CBF = 0.79 mL/g/min; Fig. 3, panels I, L, and O) stalls within R ≤ 150 µm were tolerated before oxygen at the center of the stall dropped to hypoxic levels (pO_2_ < 10 mmHg) (Fig. 3, panels T, U). For high local oxygen demand, local CMRO_2_ = 3.54 µmol/g/min Fig. 3, panels J, M, and P) stalls were tolerated up to R ≤ 100 µm (Fig. 3, panels T, U).

Decreasing the systemic hematocrit and arterial oxygen saturation uniformly lowered oxygen tension throughout the tissue, which is similar to the effect of increasing local CMRO_2_ (Fig. 3, panel U). Fig. 3 panel V shows the volume of hypoxic tissue in the 1×1×0.8mm^3^ in-vivo network core under nominal systemic hematocrit and arterial oxygen saturation values with respect to stalling radius and local CMRO_2_. The volume of hypoxic tissue rises to a significant level in tandem with the radius of the stalling zone once the critical stalling radius has been breached. The same analysis was also performed in a 3×3×1.2 mm^3^ synthetic VAN and similar trends were observed, as shown in Fig. S4 of the supplemental material.

### Capillary stalls decrease local oxygen tension with little effect on (macroscopic, observable) CBF, CMRO_2_

Local capillary stalling did not substantially alter mesoscale perfusion (CBF) nor oxygen consumption (CMRO_2_) for the entire domain. Overall CBF and CMRO_2_ for the whole computational domain (in-vivo, 3×3×1 mm^3^) remained nearly constant; with CBF dropping only from 0.79 to 0.78 mL/g/min and CMRO_2_ remaining constant at 2.1 µm/g/min to the first decimal place, despite stalling in a radius of up to 300 µm. Stalls had a rather small effect even when focusing only on the region of interest (core of 1×1×0.8 mm^3^); the CBF dropped from 6.26 to 6mL/g/min, oxygen turnover decreased from 2.45 to 2.11 µmol/g/min. It is worth noting that the value of CBF at the local scale (within the 1×1×0.8 mm^3^ core) is much larger than the corresponding number taken at the scale of the whole network. This illustrates that CBF (=total flow over volume) acquired at the voxel scale is not a good measure for perfusion impairment at the capillary event scale. Guibert et. al.^27^ previously observed that simply dividing blood flow by voxel volume introduces a severe scale bias in the computation of perfusion.

### Maximum tolerable local oxygen demand depends on CBF

Parametric dependence of hypoxia as a function of neuronal demand and perfusion are explored within small regions (eight ROI with cube size = 0.15 mm^3^). Parametric oxygen demand changes were simulated by modifying the Michaelis-Menten reaction rate constant M_0_ in equation (2); local blood perfusion (lCBF) was varied by suitable resistance adaptations of all segments within the region of interest. Results indicate that the maximum tolerable local metabolic demand (lCMRO_2, crit_) increases with higher perfusion. Parametric trends of critical lCMRO_2_ versus lCBF values are shown in supplemental Fig. S5, panels A-C. Variations are due to local capillary network configurations in different areas; for example, starting from nominal lCBF and lCMRO_2_ values of 0.8 mL/g/min and 2.5 µmol/g/min respectively, we found ROI capable of tolerating an up to eight-fold increase in lCMRO_2_ (up to 20 µmol/g/min in an 0.15 mm^3^ cubic region) before hypoxic tissue formed. Supplemental Fig. S5, panels D-F also illustrate the effects of blood flow and oxygen metabolism on tissue oxygenation. The transition from panels D to E depicts the formation of a hypoxic pocket (blue) after a sudden spike in oxygen turnover (R_1_ to R_2_ in panel B). A return to normal pO_2_ level (recovery from hypoxic state in E) is achieved with a higher lCBF value, as in panel F (normoxia; R_2_ to R_3_ in panel B). These results quantify the relative effects of blood flow micro-regulation on oxygen tension, a local control action achievable by vessel dilation.

### Resilience to hypoxia by increased reaction rate is greatest in regions of high vessel density

For constant lCBF inside the cubic region of interest, mean tissue oxygen tension 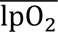 follows a trend 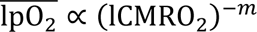 with the slope of the regression equal to the exponent −*m* in a logarithmic plot, (Fig. 3, panels E-G), which can serve as a measure of tissue resilience to a surge in metabolic demand. When the slope is flat, the mean oxygen concentration changes little with demand indicating higher resilience to O_2_ drops under neuronal demand fluctuations.

Based on this definition, tissue regions with high resilience (i.e. flat slope) can be systematically detected and the extravascular space divided into two transport regimes: (i) transport and (ii) terminal/reactive zone. We found the highest resilience near major vessels due to the oxygen reserve by blood convection and refer to these domains as the *transport zone* (Fig. 3, panel G). On the other hand, Panels E and G of Fig. 3 show regions of the distal t*erminal/reactive zone*. Here, tissue locally has moderate to high resilience depending on capillary density, with the farthest regions exhibiting a direct correlation between distance and concentration. Accordingly, capillary density ensures sufficient oxygen availability; with high capillary density needed for robustness.

In zones occupied by very thin capillaries (*terminal/reactive* zone), local perfusion (lCBF) poses a limit on oxygen availability because slender vessels may be fully depleted (lOEF=100%). By contrast, thick vessels like penetrating arterioles supply not merely their local neighborhood but in addition service downstream tissue. It follows that the lCBF of a region with large vessels exceeds levels required for local oxygen demand, because most of the blood merely passes through the local domain to irrigate downstream portions of the vascular hierarchy. An lCBF reduction or lCMRO_2_ increase in the *transport* zone (region containing a penetrating arteriole, see Fig. 3, panel G) can be easily tolerated, but can cause hypoxic damage in the terminal/reactive zone (mainly occupied by capillaries, see Fig. 3, panels E and F).

### Microscale flux analysis. Oxygen flux drops rapidly with distance to nearest vessel

We performed microscale flux analysis to quantify the protective mechanisms that bestow resilience. The dimensionless quantity *ρ* for each tissue voxel is defined as the ratio of extravasating diffusive O_2_ flux to its metabolic turnover (uptake); a quantity related to the well-known Damköhler number.^28^ Accordingly, high values of *ρ* indicate a higher rate of diffusion compared to reaction, while the converse signifies dominant consumption. Data show diffusive fluxes dominate near blood vessels (*ρ* high). The length of O_2_ flux vectors in panels B and C of Fig. 4 rapidly drop with distance, *d*, from penetrating arterioles (B) or capillaries (C). The trend captured in the inset to Fig. 4A is exponential *ρ̅* ∝ *d*^-O.44^, where *ρ̅* is the mean *ρ* value over voxels at a given distance, *d*.

**Figure 4.**
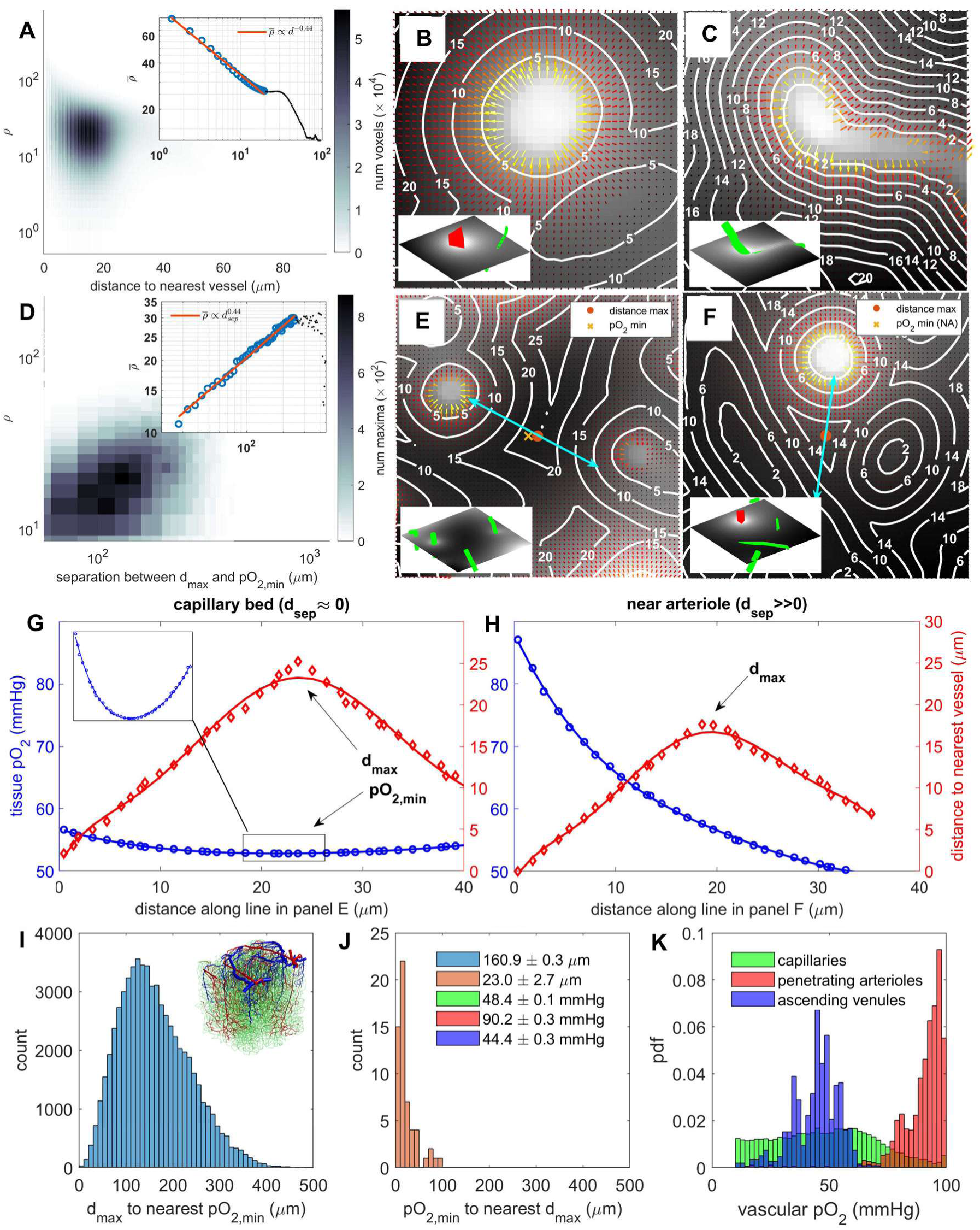
Relationship of vessel distance maxima, tissue pO_2_ minima, and the ratio of diffusive outflow to metabolic turnover, *ρ*. A) 2D histogram of flux ratio *ρ* to distance to nearest voxel *d* in each tissue voxel. Inset: mean *ρ* value, *ρ̅*, per distance value with linear fit in logarithmic scale. Black dots are values with very few voxels. B-C) Tissue pO_2_ fields near a penetrating arteriole and capillary respectively with diffusion gradient vector field and contours of nearest vessel distance. Vector field is colored according to magnitude. Inset: context view showing adjacent PA (red) and capillary (green). Voxel side length ℎ = 1.3 *μ*m. D) 2D histogram of separation *d*_sep_ (Euclidean distance) between local distance maxima *d*_max_ and local tissue oxygen minima pO_2,min_. Inset: mean *ρ* value, *ρ̅*, per separation distance value with linear fit in logarithmic scale. Black dots are values with very few voxels. E-F) Tissue pO_2_ fields and distance contours near coincident distance maxima with low and high separation values respectively. In panel F the oxygen minimum is out of view. Inset: context view showing adjacent PA (red) and capillaries (green). G-H) Values of tissue pO_2_ and nearest vessel distance along the indicated line segments in panels E and F respectively. I) Histogram of separation distance between each distance maximum and the nearest pO_2_ minimum. Inset: the in-vivo network associated with the data from this figure. J) Histogram of separation distance between each pO_2_ minimum and the nearest distance maximum. K) Estimated pdfs (normalized histograms) of vascular pO_2_ in capillaries, penetrating arterioles, and ascending venules respectively. Displayed statistics are the mean ± SEM for distance and pO_2_ measurements.

### Oxygen minima do not coincide with distance maxima near large suppliers

Krogh’s view of oxygen transport suggests that capillary distances correlate with (hypoxic) extrema in the pO_2_ field.^17^ To test this idea, we computed for each point the separation distance *d*_sep_ between the nearest local oxygen minimum and the nearest local distance maximum as in

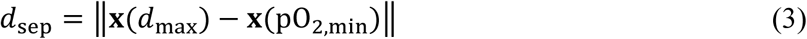

where **x**(*d*_max_) is the position of the nearest local maximum in the tissue-to-vessel distance map *d*, and **x**(pO_2,min_) marks the nearest local minimum in the tissue pO_2_ field.

In the *transport* zone (large *ρ* value, near vessels with strong transport fluxes relative to metabolic reactions), local distance maxima and pO_2_ minima are widely separated, indicating that oxygen minima do not coincide with vessel distance maxima. An example of this is shown in Fig. 4 panels F and H which show pO_2_ and vessel distance near a penetrating arteriole and capillary. Steep oxygen field gradients are shaped by strong fluxes from the penetrating arterioles which overwhelm the comparatively smaller flux from the nearest capillaries. The *lack of correlation* between pO_2_ and distance in this case is clearly shown in panel H, which shows oxygen decreasing monotonically with distance from the penetrating arteriole regardless of the distance maximum. In this *transport* zone, diffusive oxygen fluxes dominate metabolic consumption rates enabling oxygen to traverse great distances from vascular sources. In the transport zone, which is typically located around large sources such as branches of the penetrating arterioles, capillaries would not be needed to ensure sufficient exchange to tissue.

In the *terminal/reactive* zone (small *ρ* value, far away from vessels with diminishing O_2_ transport relative to metabolism), oxygen concentration rapidly decays with distance from the nearest vessels. In the *terminal region only*, distance maxima and pO_2_ minima often coincide as shown in the examples of Fig. 4 panels E and G, which show pO_2_ and vessel distance near two capillaries of roughly equal oxygen partial pressure. Oxygen and distance values along the indicated line segment are shown in panel H, clearly showing closeness of oxygen minimum to distance maximum.

The correlation between dimensionless flux, *ρ̅*, and separation distance is captured in the inset to Fig. 4, panel D, which shows an exponential relationship of the form 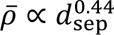 where *ρ̅* is the mean value of *ρ* over all tissue voxels with a given *d*_sep_ value (Fig. 4, panel D inset).

All simulations in this section were performed using in-vivo data from the somatosensory cortex in mouse^16^ as well as in a synthetic network.^20^ The results suggest that the separation distance between local distance maxima and oxygen minima is never zero (*d*_sep_ ≫ 0). These results contradict predictions based on the Krogh model, where *d*_sep_ is always zero. Most local distance map maxima are further away (mean difference is 161 um) from the nearest local minimum in the pO_2_ field as seen in Fig. 4, panel I. On the other hand, pO_2_ minima are necessarily always close to a distance maximum (mean d = 23 µm). **Our study suggests that a distance map is not a reliable predictor of O_2_ distribution (**Fig 5**, panels G-H).**

**Figure 5.**
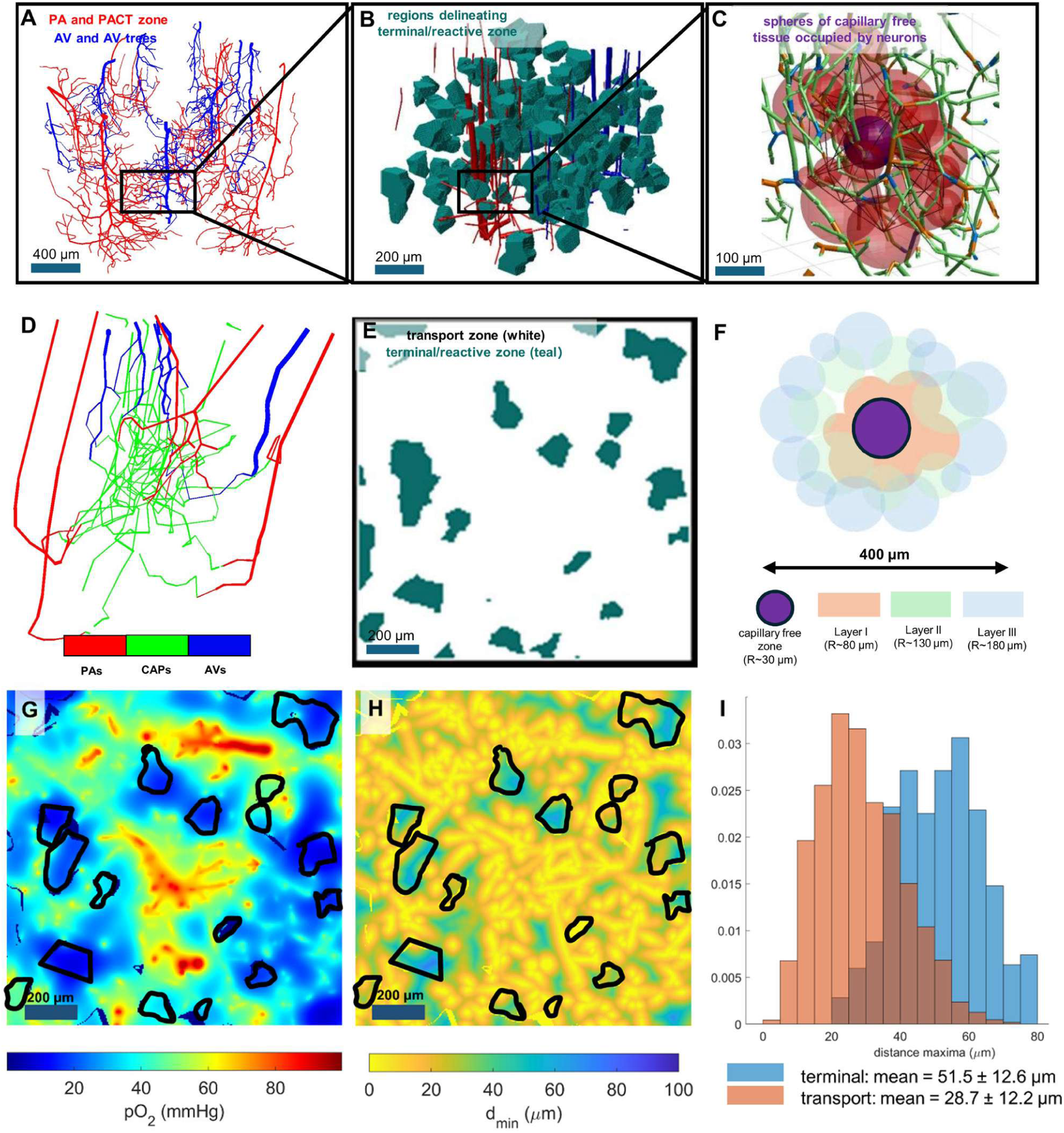
Multi-scale overview of the transport, terminal/reactive zones, and neurovascular “cages” in the microcirculation.A) Depiction of trees of penetrating arterioles (PA-red) with branching post-arteriolar capillaries belonging to the post-arteriole capillary transitional (PACT) zone^29^ and ascending venules (AV-blue). B) Blobs (teal) delineating regions of the terminal/reactive zone, encompassing tissue voxels for which d_sep_<d_max_ for the nearest local distance maximum d_max_. C) Spheres (purple) encompassing capillary free tissue (with radius equal to d_max_) and surrounding capillary segments (green; bifurcations in blue, unions in orange). D) Color coded subnetwork of penetrating arterioles (red) branching into capillaries (green) and draining into ascending venules (blue). E) Cross-section of tissue space with negative space (in teal) indicating the far terminal/reactive zone; white spaces correspond to the transport zone. F) Schematic picture of central capillary free zone (purple) surrounded by three layers of cages (layer I – pink, layer II – green, layer III – gray). In the transport zone, the mean capillary-free sphere radius (d_max_) is about 30 μm; hence, layers of capillary cages lie in approximately concentric spherical shells at radii of 60, 120, and 180 μm from the sphere center. G-H) There is dissimilarity of tissue-to-vessel distance and tissue pO_2_ maps because patterns in panel G and H do not match. G) Cross-section of tissue pO_2_ map with terminal zone overlaid in black. H) Cross-section of tissue-to-vessel distance (d_min_) map with terminal zone overlaid in black. The pO_2_ and distance maps correlate more strongly in the terminal zone, but correlate little elsewhere. I) Histograms of d_max_ values show that distance maxima are greater in the terminal/reactive zone than in the transport zone. Distance maxima and oxygen minima only coincide in regions of low capillary density in the terminal/reactive zone.

### The oxygen field is shaped by the capillary length density only in the terminal/reactive zone

Fig. 4, panel K shows the vascular oxygen partial pressure in capillaries, penetrating arterioles, and ascending venules. Penetrating arterioles tend to be highly oxygenated, hence they serve as sources and large suppliers centered in the transport zone. In addition, numerous thinner vessels (categorized as capillaries based on their small diameter) can carry highly oxygenated blood and thus may also engender a corresponding transport zone around them. These high capacity segments include the capillaries of the post-arteriole capillary transition (PACT)^29^ zone, see Fig. 5, panel A.

Because local pO_2_ minima tend to coincide with local distance maxima only in the terminal/reactive zone, it is fair to conclude that the tissue oxygen field is shaped by the capillary network only when far away from the wide-reaching branches of penetrating arterioles which release significant transport fluxes.

### The terminal/reactive zone comprises about 10% of the tissue volume

How big are the *transport* and the *terminal* zones? To obtain an unambiguous estimate of their respective volumes, we assign tissue voxels to the terminal zone if the nearest local maximum in the distance map is closer to a local minimum in pO_2_ than it is to the nearest vessel. With this definition, the *terminal/reactive* zone occupies about 10% of the total volume. Its shape conforms to a collection of approximately polyhedral blobs situated in between gaps of high-capacity vessels mainly between large branches of penetrating arteriolar trees: see Fig 5 panels B and E.

Conversely, 90% of the tissue volume falls into the *transport* zone in which the presence of penetrating arterioles and their high-capacity branches dominate the oxygen transfer inducing fairly uniform pO_2_ tension. Hemodynamically, highly conductive blood flow channels run through the transport zone and infiltrate the tissue matrix, reaching even the deepest corners located in terminal zones (Fig. 5 panel D). These capillary vessels branching from the main PA trunks (and conversely the AV main tree) have been identified independently on physiological grounds as elements of the PACT zone. **The view of the capillary bed as a homogenous, porous medium is incompatible with this finding.**

### Trends in tissue oxygen with distance from penetrating vessels are reproduced in simulations

To validate the oxygen transport model, the average arterial oxygen saturation and systemic hematocrit values measured in cohorts of young, middle-aged, and old mice were input as parameters into oxygen simulations with corresponding overall CBF and CMRO_2_ values.^2^ Observed trends in tissue oxygen tension with respect to distance from penetrating arterioles and ascending venules were faithfully reproduced in simulations: see Fig. 6.

**Figure 6.**
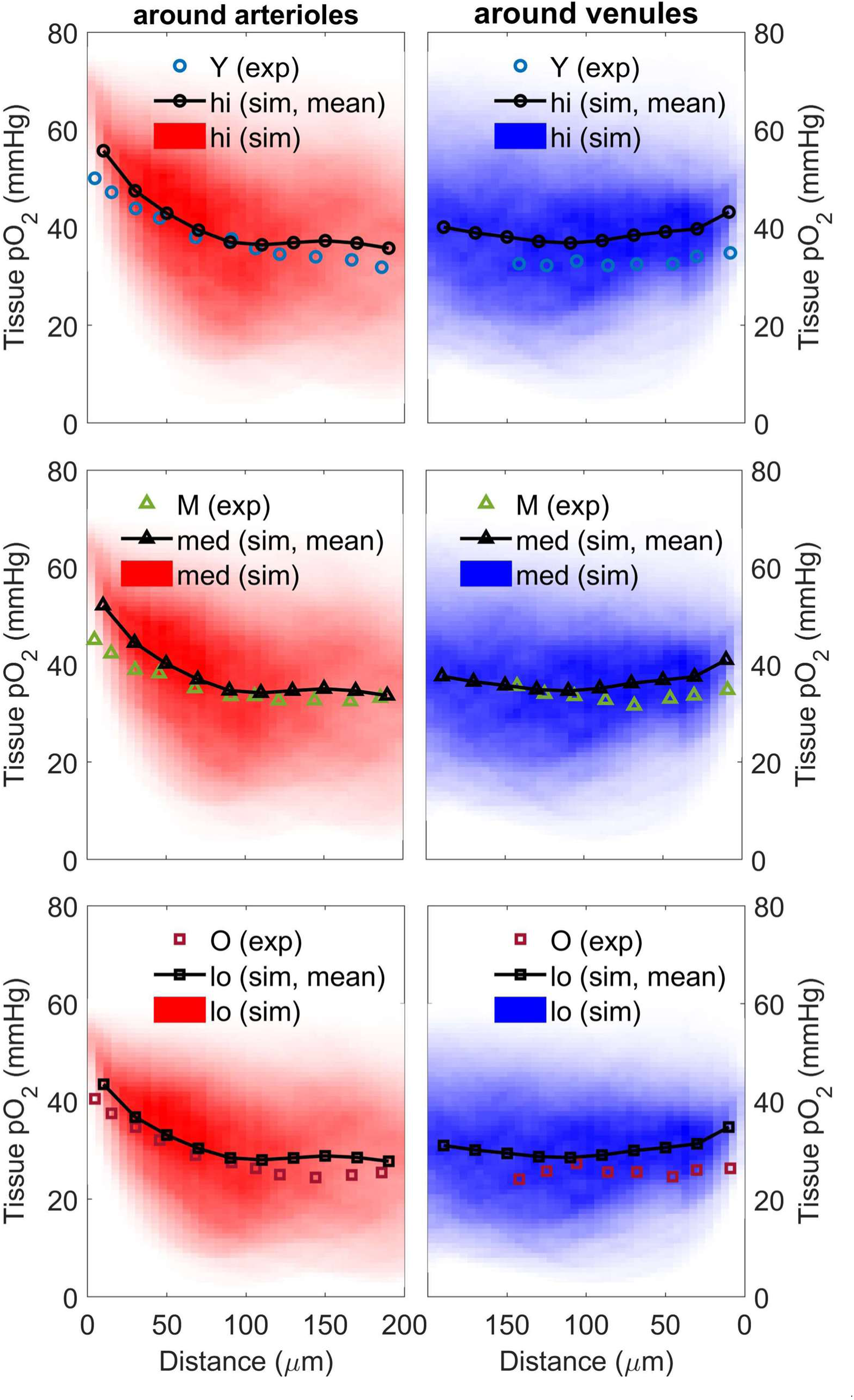
Comparison of experimental and simulated trends in tissue pO_2_ of awake young (Y), middle-aged (M), and old (O) mice with respect to distance from penetrating arterioles and ascending venules. 2D histograms of tissue pO_2_ show density of simulated tissue points. Trend-line of mean values over each 25 µm layer match well with experimental data.^2^

### The model is validated by reproducing hypoxic micro-pockets in the aged mouse brain

Next, we explored the effect of age-induced micro-stalls on oxygen perfusion of tissue. We examined conditions necessary to produce hypoxic micro-pockets that were experimentally observed in aged mouse populations in Moeini et. al.^2^ Simulations matched size and frequency of experimentally observed hypoxic pockets under conditions of reduced arterial saturation and systemic hematocrit which account for the effects of aging: see Fig. 7. The frequency of hypoxic tissue is increased further when up to 15% of capillary segments are stalled.

**Figure 7.**
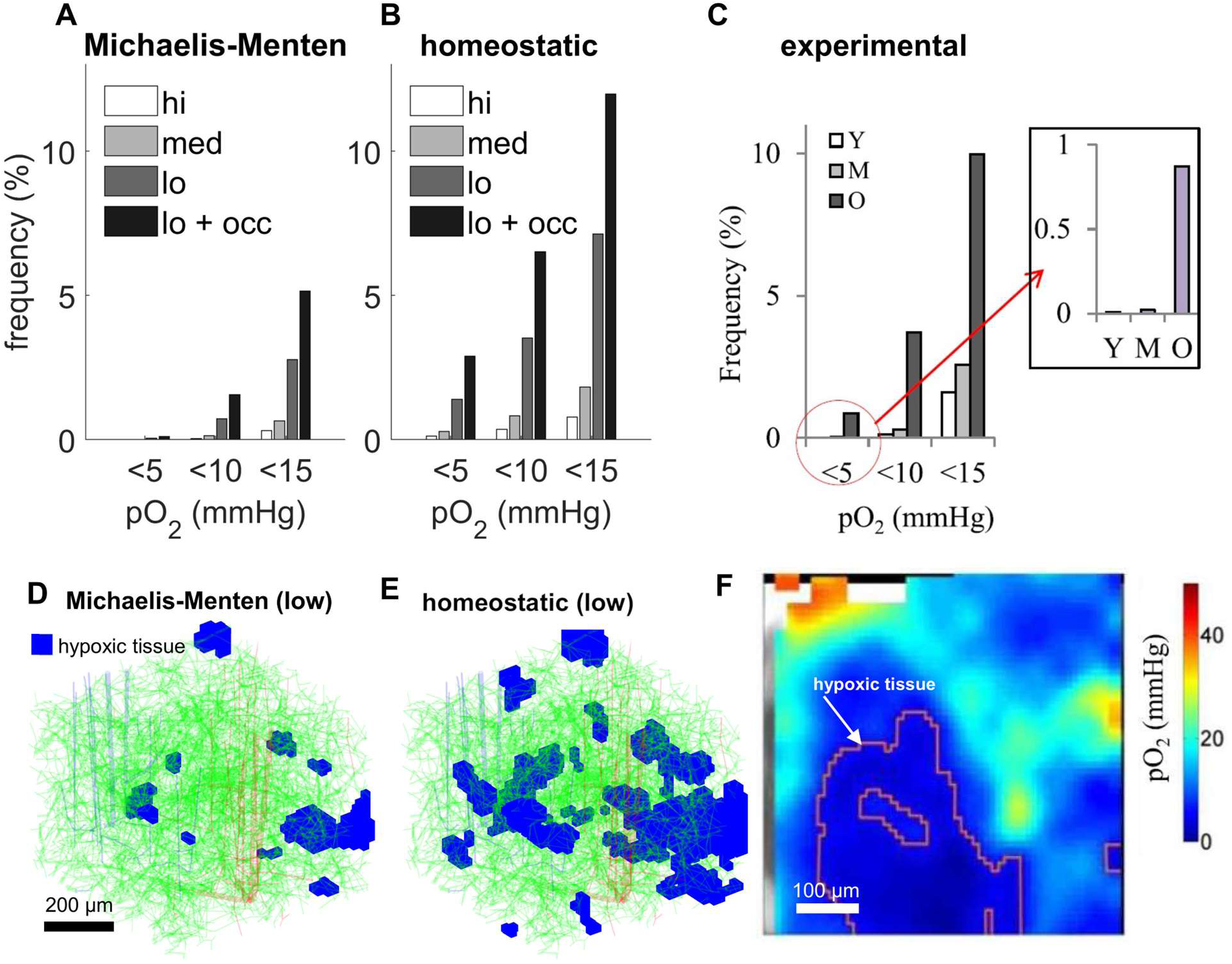
Hypoxic pockets in the young and old mouse brain. A-C) Volume of hypoxic tissue in young and old mouse brains with respect to changing inlet oxygen levels. A and B are simulated, C is from experiments in living mice.^2^ D-E) Pockets of hypoxic tissue under conditions of low oxygen supply and no capillary stalling with respect to different reaction kinetic models in the 1×1×0.8 mm^3^ core of the synthetic network. F) Experimental evidence for hypoxic pockets in awake, aged mice.^2^ Panels C and F reproduced from Moeini et. al.^2^ with the author’s permission.

To emulate the conditions of young, middle aged, and old mice, the inlet oxygen levels were modified. The high, medium, and low levels are characterized by arterial oxygen tension values of 130 mmHg, 100 mmHg, and 70 mmHg respectively, and by systemic hematocrit values of 0.5, 0.425, and 0.35 respectively.

To test the effect of different reaction kinetics, especially under more extreme conditions, we also performed simulations using a *homeostatic*^30^ (neuronal demand driven) reaction rate model in addition to the Michaelis-Menten kinetics used throughout the paper. The homeostatic reaction rate model uses a fixed neural demand driven CMRO_2_=2.44 µmol/g/min^31^ when tissue pO_2_ greater than 5 mmHg. Under extreme hypoxia (pO_2_ < 5 mmHg), the homeostatic reaction rate swaps to supply limited first order kinetics dropping to zero linearly. As expected, we found *homeostatic kinetics* to exacerbate hypoxia. On the other hand, Michaelis-Menten kinetics unrealistically throttle reaction rates when pO_2_ levels drop long before tissue pO_2_ reaches the hypoxic level, in effect reducing neuronal O_2_ metabolism and thus causing less severe hypoxia. A detailed analysis of the implications of homeostasis in neurometabolic coupling is beyond the scope of this study, but is discussed in DiNuzzo et. al.^30^

The predictive capability of our oxygen extraction model was further validated with data from mini-stroke experiments,^23^ with details given in Fig. S3 in Supplemental material.

## Discussion

Micron-scale simulation of blood and solute perfusion in anatomically complete microvascular graphs via a numerically optimized *dual-domain technique* captured the network effects of capillaries on oxygen supply to surrounding brain cells. Realistic simulation of the neurovascular coupling in the microcirculation enabled quantification of oxygen micro-gradients as functions of network configuration (capillary length density and tissue to vessel distance functions) and hemodynamics. Our study suggests that oxygen transport and metabolic processes in the capillary bed are not homogeneous, but instead occur in two separate regimes: (i) *transport* and (ii) *terminal/reactive* zones.

In the *transport* zone occupying about 90% of the tissue volume centered around high flux vessels (Fig. 5, panels E and G) the oxygen field is dominated by large efflux due to steep O_2_ gradients, with little contribution from capillaries. In this zone, capillary density has little impact on tissue oxygenation as previously observed.^32,33^ Conversely, in the *terminal/reactive* zone (Fig. 5, panels B and E) which occupies only 10% of the volume, oxygen tension decays with distance from capillaries (Fig. 5, panels G and H). Maintenance of homeostasis therefore requires dense vessel spacing because tissue is sensitive to disruptions to network perfusion. This suggests that capillary occlusion or loss can easily be tolerated in the transport zone, but potentially impairs neuronal supply in the collection of polyhedral blobs shaping the terminal zone. Resilience to local increments in CMRO_2_ is highest in the *penetrating arteriole tree*, a structure composed of the penetrating arterioles (large suppliers) and the transition to the capillary mesh (Post-Arteriole Capillary Transitional, PACT, zone^29^), as well as in the densest sections of the capillary bed (Fig. 3, panels E-G).

Cortical tissue was found to be robust against hypoxia induced by changes in local lCBF and lCMRO_2_ under physiological conditions. The danger of hypoxia arose only under extreme conditions of decreased saturation and hematocrit, as well as large disruptions in blood flow (arresting capillary blood flow in a sphere at least 150 µm radius, or local CBF reduction, or hypoperfusion subsequent to stroke). Our experimental observations in aged mice revealed distinct hypoxic micro-pockets, and the theoretical model accurately reproduced their size and frequency. This validation—achieved without any parameter adjustments—demonstrates the robustness and predictive power of the network-based simulation in capturing microvascular resilience to oxygen supply deficits. Our study did not attempt to generate histological quantification of damaged tissue. The degree of network resilience and re-distribution of flow paths after stalling was predicted by our mechanistic approach without parameter adjustments; our stationary analysis did not explore possible control counteraction to avert hypoxia.

Our observations do not support the notion that cortical solute exchange can be abstracted as *passive transport* in homogenous porous media. Our study also cautions against the practice of performing microcirculatory transport analyses on isolated capillary lattices stripped of their *penetrating arteriolar trees* or segments of the ascending venous trees. Solute exchange processes are heterogeneous and qualitatively different in the *transport* and *reactive* zones.

Our study underscores the significant role of the *penetrating arteriole tree* that includes the PACT zone in regulating flow within the capillary bed.^29,34–36^ Low-order branches in the capillary hierarchy, which emanate from the main trunk of penetrating arterioles (PAs), exhibit high conductance and carry oxygen-rich blood. Together with main trunk of PAs, they constitute the backbone of the transport zone. PACT segments act as gateways to capillary perfusion, identifiable through hydraulic network connectivity—even without biomarkers. Dual support by hemodynamic analysis here and prior histological pericyte localization^29,34,35^ substantiates its role in regulating microcirculatory blood flow. Our study also suggests that the location of thin-stranded pericytes primarily in the downstream reactive zone, is ideally positioned to detect subtle oxygen tension changes during activation.

Capillary distance maps do not match oxygen tension level sets; especially in pathologies, where network effects play an important role. Our study shows that local distance maxima and local pO_2_ minima generally do not coincide except in the *terminal zone* (occupying only 10% of the space in locations far removed from penetrating arterioles). The terminal zone is more vulnerable to vessel rarefication, so that a supply limitation in extreme pathological conditions (e.g. hypoperfusion) may fail to meet tissue demand thus potentially impairing neuronal function. In these regions which we succeeded in delineating as collections of convex polyhedra occupying isolated spaces between PA and AV trees, hypoxic conditions will occur first and most readily. However, O_2_ supply under physiological conditions was shown to be very robust due to network density and high-capacity perfusion in the transport zone, tolerating both large drops of CBF as well as rarefaction of capillaries (stalling zones) of up to 150 µm radius without causing hypoxia.

Simulations of blood flow regulation using quasi-dynamic neuronal activation under homeostatic transitions showed that CBF, CMRO₂, and BOLD signal trends aligned with a those suggested by Buxton (2010),^37^ with more details given in supplement section 9. Notably, average tissue pO₂ throughout a 1 mm^3^ volume remained nearly constant during activation, suggesting that oxygen supply is not a limiting factor under normal physiological conditions. Moreover, our quasi-dynamic simulations of neuronal activation demonstrate our ability to reconstruct the BOLD signal from first principles. The current work focuses on elucidating the stationary conditions under which persistent hypoxic pockets can exist, and the role of the network structure on shaping the tissue oxygen field. Future work will aim to compute the unsteady coupled blood-flow and oxygen fields in order to include mechanisms of dynamic control.

It is worthwhile to further contextualize organizational principles of the microcirculatory oxygen supply. The illustration of Fig. 5 panel C generated by zooming into a small section of the somatosensory cortex delineates neuronal tissue free of capillaries by colored spheres. We found that each capillary-free sphere is on average surrounded by one capillary bifurcation and its three connected vascular segments. We posit that the size of the capillary free space, we term it a *<u>cage</u>*, is determined by the metabolic demand of the embedded neuronal and glial cells in normal signaling and especially during activation, which cannot exceed the capacitance of supply by blood vessels surrounding each sphere. The tissue is resilient against stalling of one bifurcation, or even bifurcations of adjacent *cages*. Stalling in blood vessels located inside a region corresponding to up to two layers of sphere packings can be tolerated (Fig. 5, panels F and I). Hypoxia occurs when all capillaries up to a radial distance of about 150 µm are occluded, which corresponds to two to three concentric layers of capillary-free zones.

### Limitations

Our analysis focused on cortical blood supply; possible extensions to deep regions of the brain or white matter would further enhance the significance of mechanistic models for stroke, aging and small vessel disease (SVD). Results presented here for rodents do not translate directly to humans due to differences in collateralization as well as possible changes in the architecture of the pial arterial blood supply.^38^ These issues are beyond the scope of this paper. Experimental validation was limited to data collected in male mice; despite this, the framework could help elucidate differences between sexes.

### Conclusions

The dual-domain method enabled high resolution simulation of oxygen flux distribution at micron level resolution to determine the origin and degree of tissue resilience to hypoxia against local flow occlusion or surges in metabolic oxygen demand. Direct mechanistic simulation in realistic anatomy may be a promising approach towards understanding neurovascular and neurometabolic coupling principles underpinning the BOLD signal in functional neuroimaging.

Results provide support for the notion that oxygen extraction is not a limiting factor in neuronal oxygen supply under physiological conditions. Thus, oxygen supply does not seem to be a limiting factor in resting nor in activated states. Moreover, network configuration and overcapacity of blood supply lend a robust degree of microvascular reserve against hypoxia.

Hypoxic conditions were reached only in the presence of pathological deficits in systemic hematocrit and arterial oxygen saturation as might occur in aging, or very low focal CBF under conditions of widespread capillary stalls or stroke induced hypoperfusion.

## Acknowledgement.

The authors gratefully acknowledge Drs. Xiang Ji and David Kleinfeld of UC San Diego for the use of their whole mouse vasculature data set.

## Author contribution statement

All authors have contributed and read the manuscript.

Thomas Ventimiglia: Conceptualization, Data Analysis, Theory, Software, Composition Frédéric Lesage: Conceptualization, Imaging, Data Analysis, Supervision, Composition Andreas A Linninger: Conceptualization, Theory, Supervision, Composition

## Supplementary Information

Nine expository sections, six figures, and one table are provided as supplementary information.

