## Supplemental Information for "Microvascular network organization and hemodynamic perfusion protect the brain against hypoxia"

### Supplemental Material

1. **Construction of dual-domain graph.**

Each edge of the network has a pair of endpoints $\mathbf{p}_{1}$ and $\mathbf{p}_{2}$ whose 3D Cartesian coordinates are known, and a diameter $d$. Denote the Euclidean length of the edge by $\mathcal{l=}\left\| \mathbf{p}_{2}-\mathbf{p}_{1} \right\|$. Construct a right-circular cylinder of length $\mathcal{l}$ and diameter $d$ with hemispherical endcaps whose centerline is placed at the origin and is vertically aligned with the positive $z$-axis; that is, the centerline of the cylinder is the line segment connecting the points $(0,0,0)$ and $(0,0,\mathcal{l)}$. Call this shape a capsule. We sample some points from the surface of the capsule and label them as surface points, then construct more concentric capsules in its interior and label them as interior points. We then construct the 3D rotation matrix (1) which rotates the vertical centerline of the capsule (the vector $(0,0,\mathcal{l)}$) onto the vector $\mathbf{p}_{2}-\mathbf{p}_{1}$, apply this rotation to all interior and surface sample points of the capsule, and translate them into the position of the edge (if the $3\times3$ rotation matrix is $R$ and the three-row matrix $P$ has the coordinates of the center points of each voxel as its columns, then the matrix product $RP$ has the coordinates of the rotated center points as its columns. We then add the vector $\mathbf{p}_{1}$ to each column to translate them into place). Let $\mathbf{b}=\frac{\mathbf{p}_{2}-\mathbf{p}_{1}}{\left\| \mathbf{p}_{2}-\mathbf{p}_{1} \right\|}$ denote the unit vector parallel to the centerline of a given vessel. If $b_{x}^{2}+b_{y}^{2}\neq0$, then the rotation matrix $R$ can be derived from the Euler-Rodrigues formula^1^ as:

$$R=\left( \begin{matrix} 1 & 0 & b_{x} \\ 0 & 1 & b_{y} \\ -b_{x} & -b_{y} & 1 \end{matrix} \right)+\frac{b_{z}-1}{b_{x}^{2}+b_{y}^{2}}\left( \begin{matrix} b_{x}^{2} & b_{x}b_{y} & 0 \\ b_{x}b_{y} & b_{y}^{2} & 0 \\ 0 & 0 & b_{x}^{2}+b_{y}^{2} \end{matrix} \right)$$

(1)

Otherwise, if $\mathbf{b}=(0,0,1)$ then $R$ is an identity matrix, and if $\mathbf{b}=(0,0,-1)$ then $R=\left( \begin{matrix} 0 & 0 & 0 \\ 0 & 0 & 0 \\ 0 & 0 & -1 \end{matrix} \right)$.

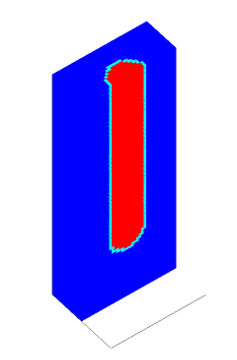
In this way we obtain the Cartesian coordinates of a correctly oriented and labelled capsule around the given edge. We then register these points to voxels in the tissue domain by determining which voxel contains each point and classify the voxels as interior lumen or endothelial wall according to the labels of the points: see Fig. S1. In the case that two points with different labels lie in the same voxel, we identify the voxel as belonging to the vessel wall, as we reason that having a thick wall is preferable to having a hole in the wall. Furthermore, in the case that the edge (=vessel segment) has a diameter much smaller than the side length of the voxels, such that the entire capsule is contained within a single voxel, that voxel is identified as belonging to the vessel wall. The number of sample points on the original capsule is chosen large enough so that each voxel contains at least one of the rotated and translated capsule points, this way we avoid holes in the vessel walls.

Fig. S1. Cross sections of a voxelized capsule. Red voxels are tagged as interior lumen, blue as extravascular tissue, and cyan as endothelial wall.

For each edge we follow the procedure in the preceding paragraph to determine the voxels comprising the surface and interior of its capsule. Then for each node of the network we determine the edges incident to it, take the halves of their capsules closest to the node, identify their interior and surface voxels and eliminate the overlap. In this way we associate to each node a collection of voxels representing the surface of its associated vascular control volume; a conjunction of half-capsules. In total, we have a voxelized representation of the vascular network, as in Hartung et. al.^2^ On the scale of the large vessels, such as the penetrating arterioles, a vascular node is associated with multiple voxels (=tissue nodes): those coinciding with the interior and surface of its control volume. Conversely, on the scale of the smallest vessels, such as the capillaries, a vascular control volume might be so small as to fit entirely in the interior of a single voxel. In that case, the node is associated with one voxel only, and that voxel is labelled as belonging to the vessel wall. In this case there may be multiple nodes associated with a single voxel.

The initially separate vascular and tissue graphs are coupled into a single dual-domain graph by including an edge connecting each node to the tissue voxels associated with the surface of its control volume. It follows that are coupled into a single dual-domain graph by including an edge connecting each node to the tissue voxels associated with the surface of its control volume. It follows that diffusive solute transport within and between the coupled domains is captured by the weighted graph Laplacian of the entire coupled graph, as in equation (1) of the main document.

2. **Simulation of hemodynamics, plasma-skimming, and dissociation kinetics**

Biphasic blood flow is approximated using parabolic velocity profile assumptions on a one-dimensional vascular graph network with shear rate (diameter) and hematocrit dependent viscosity.^3–5^ The highly nonlinear and coupled bulk flow and hematocrit field computations are solved by a fixed point iteration scheme which alternates between solving two linear subsystems. These iterate between computing a hematocrit field by solving a linear convection problem in which RBC flux is adjusted by uneven splitting, and computing steady blood-flow with viscosity field derived from the current iterate of the hematocrit field. The blood-flow and hematocrit fields typically converge within a handful of iterations of these two alternating solution procedures.

*Flow and pressure (f,p) simulation for given hematocrit field, h (linear elliptic problem).*

Flow **f** and hematocrit **h** vectors have components indexed by the edges of the vascular graph, and pressure vector **p** is defined on the nodes of the vascular graph. Flow on an edge is related to pressure drop by the Hagen-Poiseuille relation $\alpha\left( d,h \right)\cdot f=\Delta p$, where the resistance $\alpha$ is a nonlinear function of the diameter $d$ and hematocrit $h$ of an edge.^4,6^ Furthermore, mass balance requires that at every node the sum of flows on entering edges equals the sum of flows on leaving edges. Given Dirichlet boundary conditions for pressure at the inflow and outflow nodes and a known hematocrit field **h** (initially set to be uniformly equal to the systemic hematocrit), the system of equations to solve for the unknown flow **f** and pressure **p** vectors simultaneously is given by the matrix equation (2):

$$\left[ \begin{matrix} \mathbf{A}\left( \mathbf{h} \right) & -\mathbf{C}_{1}^{T} \\ \left( \mathbf{I-D} \right)\mathbf{C}_{1} & \mathbf{D} \end{matrix} \right]\left[ \begin{matrix} \mathbf{f} \\ \mathbf{p} \end{matrix} \right]=\left[ \begin{matrix} 0 \\ \mathbf{D}\bar{\mathbf{p}} \end{matrix} \right]$$

(2)

Here, **A(h)** is a diagonal matrix of edge resistances dependent on the current hematocrit field, **C**_1_ is the directed incidence matrix of the vascular graph:

$$\left[ \mathbf{C}_{1} \right]_{ij}=\left\{ \begin{matrix} 1 & if edge j leaves node i \\ -1 & if edge j enters node i \\ 0 & \mathrm{otherwise} \end{matrix} \right.,$$

(3)

**D** is a diagonal decision matrix with ones on the main diagonal for every node with a pressure boundary condition and zeros everywhere else, **I** is an identity matrix, and $\bar{\mathbf{p}}$ is a vector containing the boundary values.

We note that flow values computed in this manner may be negative because the orientation of the vascular graph is initially arbitrary. By convention, we reverse the orientation of edges so that flow values are positive where necessary.

*Hematocrit update for given hematocrit field, flow, and bifurcation geometry in the network (linear hyperbolic problem with bifurcation adjusted virtual flows θf and skimming coefficient).*

Using computed flows **f**, the segment hematocrit $\mathbf{h}$ is updated by solving the linear convection problem in equation (8), in which *virtual* geometry θ-adjusted fluxes driving RBC convection leading to uneven splits in bifurcations (as a function of diameter ratios and the drift parameter, *m*). The unknown $\mathbf{h}^{\boldsymbol{*}}$ is an auxiliary quantity representing *virtual* hematocrit at each node; more details are provided in prior works.^2–4^

Conservation of RBCs at each bifurcation is written as

$$f_{p}H_{p}=f_{1}H_{1}+f_{2}H_{2}$$

$$f_{p}H_{p}=\left( \hat{f}_{1}+\hat{f}_{2} \right)H^{*}$$

(4)

for parent vessel *p* and daughter vessels 1 and 2, and virtual hematocrit $H^{*}$ at the bifurcation node. The virtual hematocrit $H^{*}$ is defined to satisfy

$$H_{1}=\theta_{1}H^{*}, H_{2}=\theta_{2}H^{*}$$

(5)

for the kinematic plasma skimming coefficients

$$\theta_{j}=\left( \frac{d_{j}}{d_{p}} \right)^{2/m},$$

(6)

where $d_{j}$ is the diameter of edge *j*, $d_{p}$ is the diameter of its parent vessel, and the drift parameter *m*=5.25. The bifurcation geometry adjusted flows thereby satisfy

$$\hat{f}_{j}=f_{j}\theta_{j}$$

(7)

All together, the equations (4)-(7) define the system of equations (8) in the unknown vectors **h** and **h^*^** for hematocrit in every edge, and virtual hematocrit in every node:

$$\left[ \begin{matrix} \mathbf{I} & -\boldsymbol{\Phi}\left⟦ \mathbf{C}_{1}^{T} \right⟧ \\ 0 & \mathbf{M}(\hat{\mathbf{f}}) \end{matrix} \right]\left[ \begin{matrix} \mathbf{h} \\ \mathbf{h}^{*} \end{matrix} \right]=\left[ \begin{matrix} 0 \\ \mathbf{Q}_{\mathrm{in}}\boldsymbol{(}\hat{\mathbf{f}})\bar{\mathbf{h}} \end{matrix} \right]$$

(8)

Here, **Φ** is the diagonal matrix of *θ* values for each edge and $\bar{\mathbf{h}}$ is a vector of prescribed systemic hematocrit values at the inflow boundaries. The convection matrices **M**(**f**) and **Q**_in_(**f**) are

$$\mathbf{M}\left( \mathbf{f} \right)\mathbf{=-}\mathbf{C}_{1}\mathrm{diag}\left( \mathbf{f} \right)\left⟦ \mathbf{C}_{1}^{T} \right⟧\boldsymbol{+}\mathrm{diag}\left( \mathbf{D}_{\mathrm{out}}\mathbf{C}_{1}\mathbf{f} \right)$$

$$\mathbf{Q}_{\mathrm{in}}\left( \mathbf{f} \right)\mathbf{=}\mathrm{diag}\left( \mathbf{D}_{\mathrm{in}}\mathbf{C}_{1}\mathbf{f} \right)$$

(9)

where the double brackets $\left⟦ \cdot\right⟧$ denote the *positive part* operator: $\left⟦ \mathbf{X} \right⟧_{ij}=\max\left( \left[ \mathbf{X} \right]_{ij},0 \right)$, $diag(\mathbf{v})$ denotes the diagonal matrix with the values of the vector **v** on the main diagonal, and the inflow and outflow decision matrices **D**_in_ and **D**_out_ have ones on the main diagonal at the inflow and outflow boundary nodes respectively, and zeros elsewhere. More details on the construction of these matrices as a finite volume discretization of the hyperbolic solute convection problem on a graph are given in prior work.^7^

The entire two step fixed-point iteration scheme alternates between solving equations (2) and (8) until flow, **f**, and hematocrit, **h** fields converge, which is typically achieved in five to seven iterations for any size network: see Fig. S2. Note that our virtual ‘*convective’* hematocrit updates operate on a directed acyclic graph (DAG) which is guaranteed to have a solution. Our implementation of hematocrit dependent flow rate and plasma skimming overcomes convergence problems and instabilities introduced by the highly non-linear splitting rules.^8^

*Dissociation kinetics in blood.*

O_2_-dissociation from HbO_2_ bound to free (unbound) O_2_ in plasma is modeled via the non-linear Hill equation.^9^ The concentration of bound oxygen, c_v_, at a graph node is related to that of free oxygen according to a the inverting the Hill equilibrium relation:

$$K\left( c_{v} \right)=\alpha P_{50}\left( \frac{c_{v}}{C_{0}-c_{v}} \right)^{1/n}$$

(10)

where α is the solubility of oxygen in tissue, P_50_ is the partial pressure of oxygen in blood at 50% hemoglobin saturation, C_0_ is the concentration of bound oxygen at 100% hemoglobin saturation, and, n, is the Hill exponent. Parameter values are given in Table S1.

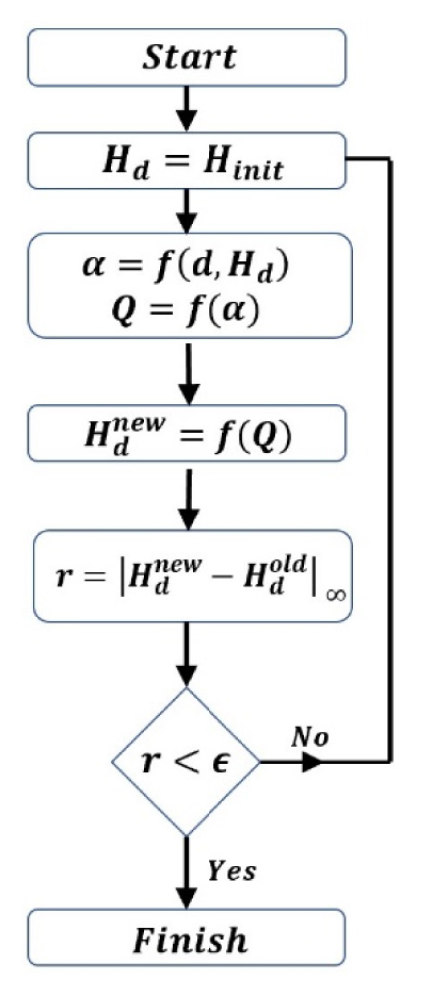

Fig S2. Flow chart describing fixed-point iteration scheme to solve for bulk blood flow and hematocrit. Reproduced from Jamshidi et. al.^4^ with the authors permission.

3. **Newton iteration**

Equation (1) of the main document is nonlinear due to Hill dissociation *K* and Michaelis-Menten reaction rates *R*; hence it is solved iteratively by Newton’s method. In each Newton step, we eliminate the vascular states **c**_v_ from the linearized equations by an equivalent Schur complement.^10^ The Schur complement implicitly embeds convective transport in blood as source terms into the reactive/diffusion processes in tissue. The reaction/diffusion equation in the 3D Cartesian tissue array with implicitly embedded convective source terms needs to also be solved iteratively due to its size.^11^ Hence, we solve for the tissue states **c**_t_ using a preconditioned Krylov subspace method (GMRES),^10^ where the preconditioning step beneficially exploits fast-sine transforms.^12^ We further improve computational efficiency by solving on a coarse grid before interpolating up to a finer grid, using the interpolated solution as an initial guess. We recover the vascular states from the solved tissue states to obtain a new Newton direction and use a simple line-search algorithm^13^ to choose a step-size that guarantees descent of residual errors. The iterates are deemed converged when the relative residual of the fully coupled system is below 10^-6^. Parameter values for the simulations are given in supplementary Table S1. Choices of nominal conditions are explained in supplement section 4. More mathematical details on the design and implementation of the solution algorithm is provided elsewhere.^7,14^

4. **Parameter values and physiological reference simulations:**

Simulation parameters are given in Table S1. We used experimental CBF, CMRO_2_, and OEF values obtained by Zhu et. al.^15^ as nominal reference conditions for simulations. The nominal CBF was calibrated by adjusting the pressure boundary conditions for blood flow to achieve the desired overall inflow to the network. The maximum metabolic rate in the Michaelis-Menten equation ($M_{0}$ in equation (3) of the main document) was chosen to yield the desired CMRO_2_ value. For simulations involving the the 3x3x1 mm^3^ synthetic network, the metabolic rate was chosen to achieve the desired CMRO_2_ in the central 1x1x0.8 mm^3^ region of interest. The systemic hematocrit and inlet arterial oxygen tension were chosen in concert with desired nominal CBF and CMRO_2_ values to obtain the desired overall OEF.

To validate the choice of nominal simulation parameters, the average arterial oxygen saturation and systemic hematocrit values measured in cohorts of young, middle-aged, and old mice by Moeni et. al.^16^ were input as parameters into oxygen simulations. These simulations faithfully reproduced observed trends in tissue oxygen tension with respect to distance from penetrating arterioles and ascending venules: see Fig. 6 in the main document.

Table S1. Physical parameters used in simulations at nominal conditions.

| **Symbol** | **Description** | **Value** | **Units** | **Reference** |
| --- | --- | --- | --- | --- |
| $H_{d}$ | Systemic hematocrit | 0.425 | NA | see text |
| $C_{0}$ | Oxygen binding capacity of hemoglobin | 0.0203 | mol/L | ^17^ |
| $n$ | Hill exponent | 2.59 | NA | ^17^ |
| $P_{50}$ | Oxygen tension at 50% hemoglobin saturation | 40.2 | mmHg | ^17^ |
| $\mathrm{apO}_{2}$ | Arterial oxygen tension | 100 | mmHg | ^16^ |
| $\alpha$ | Solubility of oxygen | 10^-6^ | mol/L/mmHg | ^18^ |
| $U$ | Permeability of vessel wall | 2400 | µm^2^/s | ^2^ |
| $\Gamma_{t}$ | Diffusivity in tissue | 1800 | µm^2^/s | ^2^ |
| $w$ | Thickness of endothelial wall | 1 | µm | ^2^ |
| $P_{0}$ | Michaelis constant; pO_2_ at half maximum metabolic rate | 10.5 | mmHg | ^17^ |
| $M_{0}$ | Maximum metabolic rate of O_2_ | 5.17$\times$10^-5^ | mol/L/s | see text |
| $\mathrm{CBF}$ | Cerebral blood flow | 0.79 | mL/g/min | ^15^ |
| $\mathrm{CMR}O_{2}$ | Cerebral metabolic rate of O_2_ consumption | 2.44 | µmol/g/min | ^15^ |
| $\mathrm{OEF}$ | Oxygen extraction fraction | 0.34 | NA | ^15^ |
| $\Delta p$ | pial blood pressure drop | 67 | mmHg | see text |

5. **Mouse Experiment Data**

The work in this paper focused on realistic simulations and comparison using previously acquired data from an aging cohort^16^ (Moeini et al. Sci. Rep. 8(1), 8219 (2018)). In that work, Young adult (6–9 month-old), middle-aged (13–16 month-old) and old (24–28 month-old) C57BL/6 J healthy male mice were obtained from the colony of aged mice of the Quebec Network for Aging Research (RQRV) and housed in 12-hr light-dark cycle until imaging. All mice were males and divided as follows: 7 young (average age = 8.8 ± 0.1 month-old), 6 middle-aged (average age = 15.3 ± 0.1 month-old), 7 old (average age = 27.0 ± 0.1 month-old)) used to record tissue pO_2_. No mice were excluded.

6. **Simulation of micro-strokes**

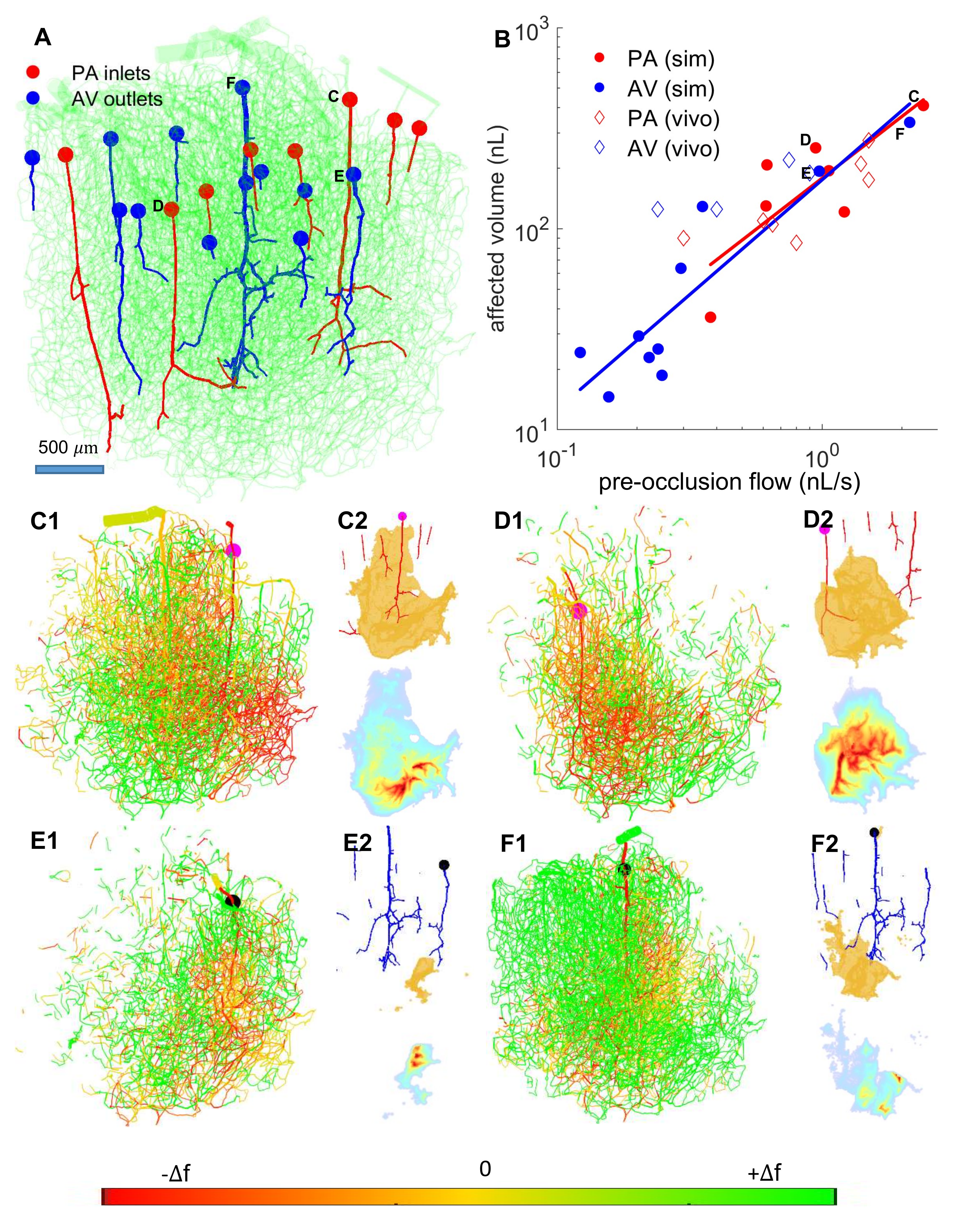

Fig. S3. Micro strokes induced by flow occlusion at the entrance to the pial surface. Outliers with volume < 1 nL associated with one penetrating arteriole and two ascending venules with very short penetration depth are excluded. A) VAN showing penetrating arterioles in red, ascending venules in blue, and all other vessels in green with low opacity. Stall locations are indicated by the circular markers at the pial surface. B) Log-log diagram of affected tissue volume versus pre-occlusion flow at the occlusion site in penetrating arterioles and ascending venules respectively. Simulated data points are plotted along with in-vivo data points from occluded PAs and AVs in the rat cortex ^19^. Solid lines represent linear regression lines for simulated data points. C-F) Effect of flow occlusion in selected penetrating arterioles (C,D) and ascending venules (E,F). Stall locations correspond to labelled markers in panels A and B. (C1-F1) Vessels with an absolute flow change of 25% or greater. Redirection of flow is indicated by flow increases in green vessels. (C2-F2) Hypoxic tissue mask (<10 mmHg pO_2_) after occlusion. Color map in lower panel indicates magnitude of relative pO_2_ in the hypoxic zone relative to nominal state. Hypoxic tissue zones C2-E2 have max diameters of $\sim600, 500,$and $150 \mu m$ respectively. Hypoxic zone in F2 is oblong, with max dimensions of $\sim250$by $750\mu m$.

7. **Stalling in synthetic VAN**

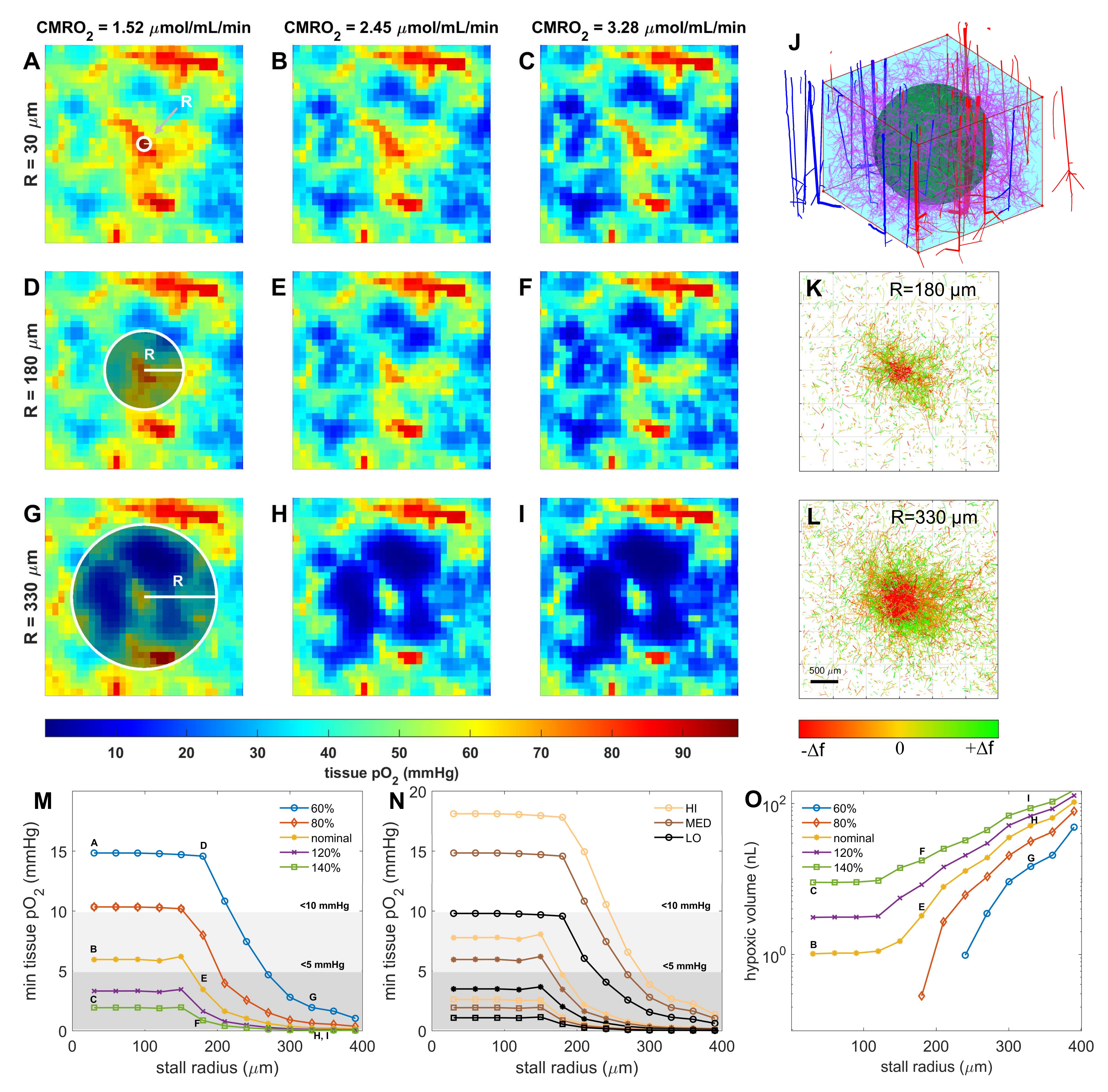

Fig. S4. The relation of capillary stalling on hypoxia in a synthetic network. A-I) Cross sections of tissue pO_2_ under different stalling and demand scenarios. Capillary stalls are confined to spherical regions of increasing radius R delineated by the green outline (displays along vertical rows). We also varied CMRO_2_ change over a central cubic ROI with dimensions 1 x 1 x 0.8 mm. (varied along horizontal columns). Hypoxia forms for large stall R and elevated CMRO_2_ indicative of network resilience of the O_2_ supply to stalling events. J) ROI and spherical stalling zone in context with surrounding network. Blue and red vessels are ascending venules and penetrating arterioles respectively. Green and magenta vessels are capillaries; only green vessels within the spherical region are stalled. K-L) Vessels with redirected flow around the central stalling zone of radius R for panels D-F and G-I respectively. Redirected flow is extensive throughout the network, far from the central focus of the stalls. M-N) Minimum tissue pO_2_ in cubic ROI and K) volume of hypoxic tissue with respect to stalling extent and reaction rate. In panels M and N, the approximate diffusion length is indicated by the intersection of each curve with the 10 mmHg isoline. The corresponding scenarios in panels A-I are indicated next to the appropriate data point in panels M and O.

8. **CBF/CMRO_2_ and hypoxia curves for 8 ROI**

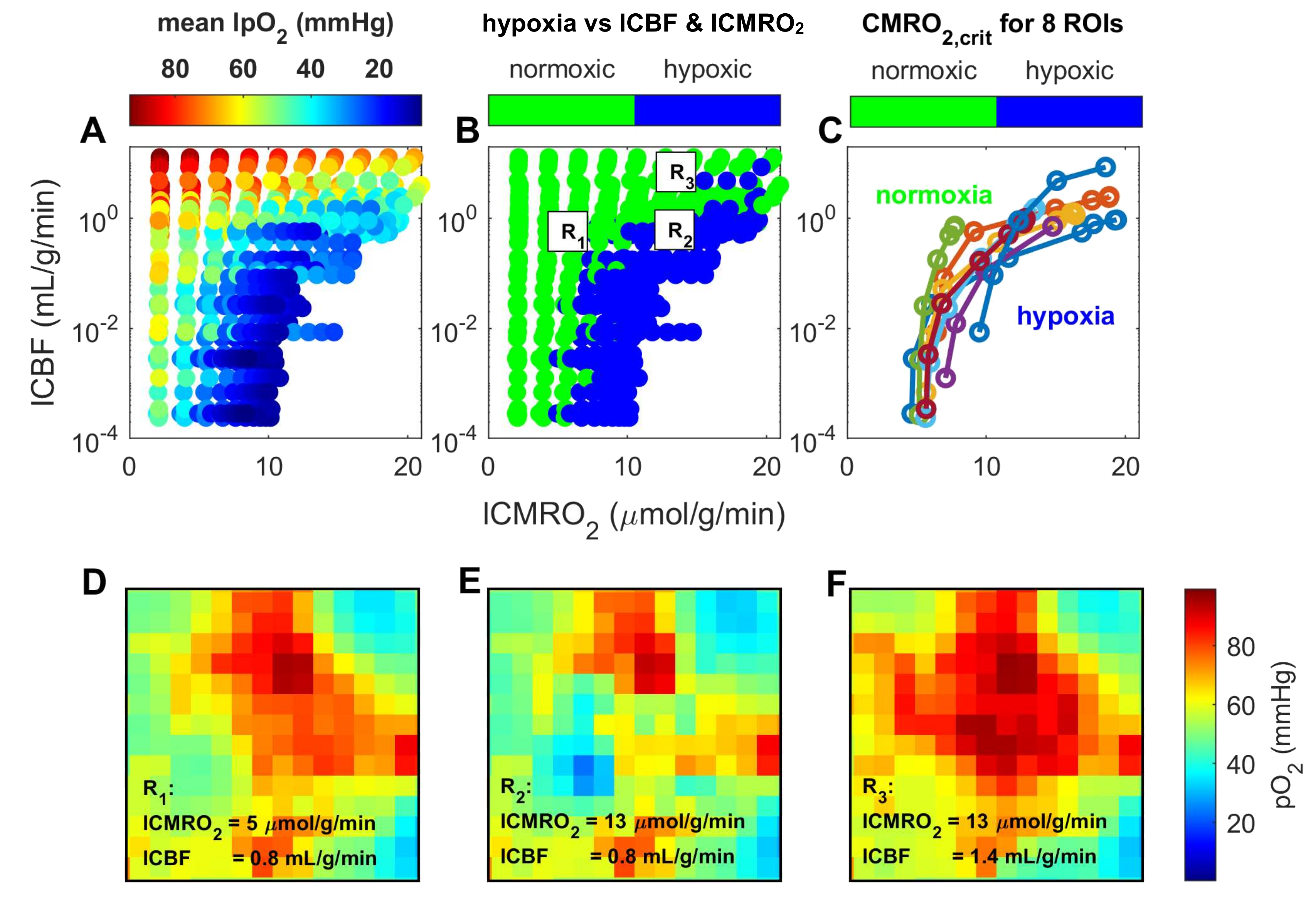

Fig. S5. Resilience of tissue ROIs to changes in local CMRO_2_ and CBF. Each ROI is a cubic voxel of side length 150 $\mu m$. A) Scatter plot of mean tissue pO_2_ in all ROIs and experiments with respect to local CMRO_2_ (increase in oxygen demand) and CBF (resistance to blood flow in vessels). B) Same as panel A with data points colored according to the presence of hypoxic tissue in the ROI. C) Critical CMRO_2_ boundary for all local CBF values. Each trend-line represents a distinct ROI. D-F) Tissue pO_2_ field in an ROI after increasing local CMRO_2_ (D to E), then after increasing CBF (E to F).

9. **Quasi-dynamic state simulation of hypothetical activation scenario**

We followed Buxton’s proposed activation scenario to compute hemodynamic and tissue state changes during activation under steady conditions. We rigorously computed blood flow in all vascular segments, blood saturation (SO_2_), oxygen tension in tissue (tpO2), and oxygen extraction fraction (OEF) with known correlation between CBF and CMRO_2_ (neurovascular coupling).We further calculated the BOLD response for the entire 1 mm^3^ voxel using Davis’ formula given in Buxton.^20,21^ Our simulations show that oxygen extraction (OEF) actually decreases and that pO_2_ is fairly constant, lending support to the notion that oxygen supply is not a limiting factor during activation. This result generates the BOLD signal from fundamental principles of blood supply and oxygen exchange.

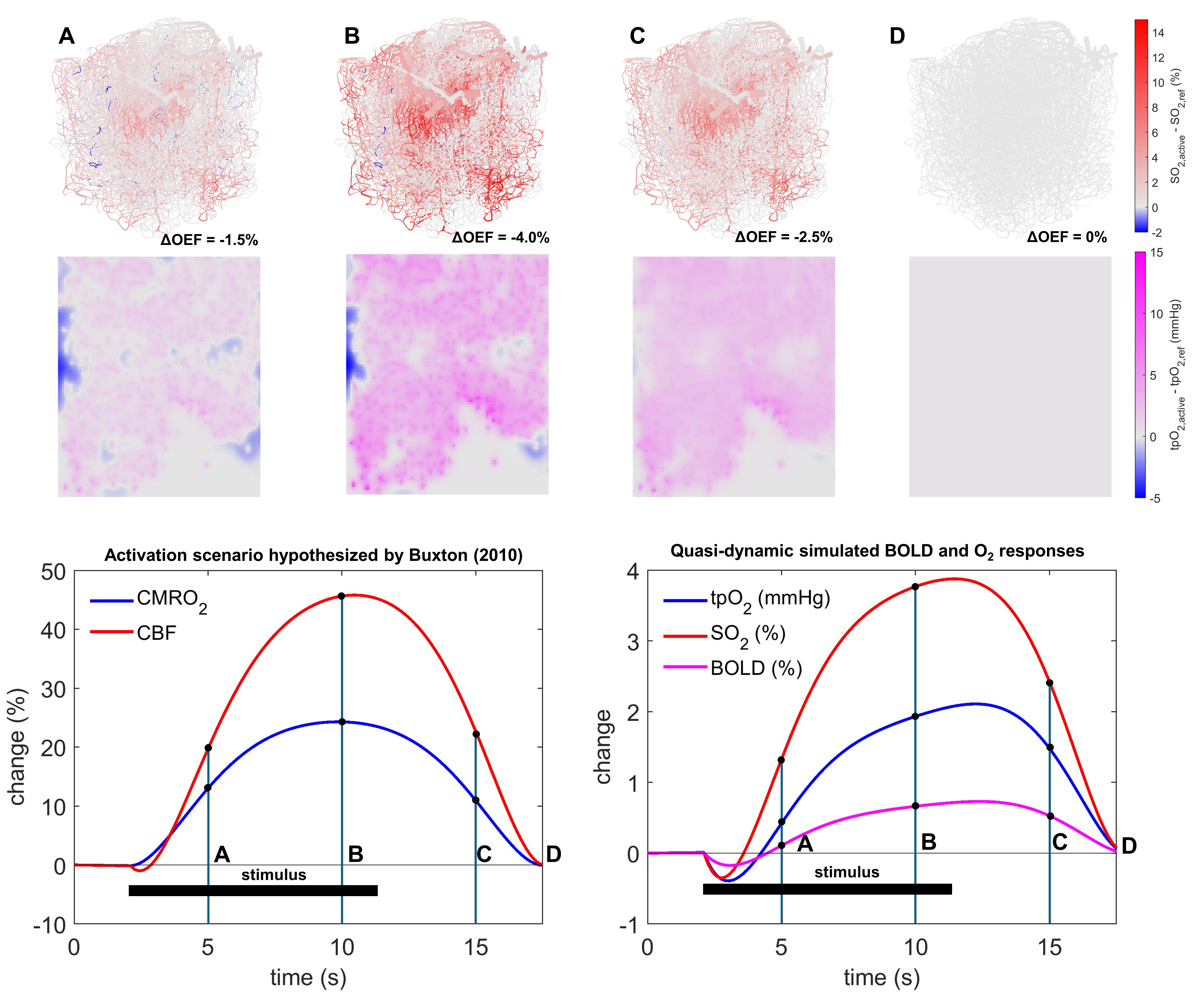

Fig. S6. Hypothetical activation scenario adapted from Buxton (2010)^20^. Panels A-D show the change in vascular pO_2_ from the nominal state in an in-vivo 1 mm^3^ vascular network^22^ after imposing the displayed changes in CBF and CMRO_2_. Bottom row (left) shows a hypothetical time-course of CBF and CMRO_2_ changes after stimulation and (right) computed changes in mean tissue pO_2_ computed by the rigorous model presented in this work, and predicted changes in the BOLD signal using an approximate model proposed by Buxton^20^. The maximum change in mean vascular and tissue pO_2_ are 3.5 mmHg and 2.0 mmHg respectively. Note that the OEF decreases with activation, implying that oxygen supply is not limiting in activation: the maximum change ΔOEF = OEF_active_ – OEF_ref_ = –4.0%.
